# Modulation of the ATP-adenosine signaling axis combined with radiotherapy facilitates anti-cancer immunity in brain metastasis

**DOI:** 10.1101/2024.09.30.615883

**Authors:** Anna Salamero-Boix, Michael Schulz, Julian Anthes, Jens Mayer, Aylin Möckl, Ioanna Tsoukala, Dominic Menger, Mohammed H. Mosa, Jenny Hetzer, Jadranka Macas, Stephanie Hehlgans, Jonas Schuck, Bastian Roller, Yvonne Reiss, Guillaume Hochart, David Bonnel, Hind Medyouf, Mariana Barcenas Rodriguez, Thomas Broggini, Marcus Czabanka, Karl H. Plate, Mathias Heikenwälder, Franz Rödel, Patrick N. Harter, Katharina Imkeller, Lisa Sevenich

**Affiliations:** Georg-Speyer-Haus, Institute for Tumor Biology and Experimental Therapy, Frankfurt am Main, Germany; Eberhards Karls University Tübingen, Department of Neurology and Interdisciplinary Neurooncology, Hertie Institute for Clinical Brain Research, Tübingen, Germany; Eberhards Karls University Tübingen, M3 Research Center for Malignome, Metabolome and Microbiome, Tübingen, Germany; Goethe University Frankfurt, Biological Sciences, Faculty 15, Frankfurt am Main, Germany; Eberhard Karls University Tübingen, Cluster of Excellence iFIT (EXC2180) ‘Image guided and Functionally Instructed Tumor Therapies’, Tübingen, Germany; Johannes Gutenberg University, Biology, Faculty 10, Mainz, Germany; Goethe University Frankfurt, Computer Science and Mathematics, Faculty 12, Frankfurt am Main, Germany; Goethe University Frankfurt, University Hospital, Institute of Neurology (Edinger Institute), Frankfurt am Main, Germany; Goethe University Frankfurt, University Hospital, Department of Radiotherapy and Oncology, Frankfurt am Main, Germany; Goethe University Frankfurt, Senckenberg Institute of Neurooncology, Frankfurt am Main, Germany; Aliri, Parc Eurasanté, Loos, France; EMBL Metabolomics Core Facility, Heidelberg, Germany; Goethe University Frankfurt, Department of Neurosurgery, Frankfurt am Main, Germany; German Cancer Research Center (DKFZ), Division of Chronic Inflammation and Cancer, Heidelberg, Germany; Ludwig-Maximilians University, Faculty of Medicine, Center for Neuropathology and Prion Research, Munich, Germany; Goethe University Frankfurt, University Cancer Center (UCT), Frankfurt am Main, Germany; Goethe University Frankfurt, Frankfurt Cancer Institute (FCI), Frankfurt am Main, Germany; German Cancer Consortium (DKTK), Partner Site Frankfurt/Mainz, Germany and German Cancer Research Center (DKFZ), Heidelberg, Germany; German Cancer Consortium (DKTK), Partner Site Munich, Germany and German Cancer Research Center (DKFZ), Heidelberg, Germany; German Cancer Consortium (DKTK), Partner Site Tübingen, Germany and German Cancer Research Center (DKFZ), Heidelberg, Germany

**Author notes:** Present address: Albert Ludwigs University Freiburg, University Hospital, Department of Neuropathology, Freiburg, Germany (Michael Schulz).

**Keywords:** tumor microenvironment, tumor immunology, immunotherapy, immunometabolism, adenosine, brain cancer, immunosuppression

## Abstract

The immunosuppressive microenvironment in the brain poses a major limitation to successful therapy for brain metastases. Here we report that blockade of the ATP-to-adenosine-converting enzymes CD39 and CD73 and the adenosine receptor A2AR in combination with radiotherapy attenuates tumor progression in a breast-to-brain metastasis model by facilitating anti-cancer immunity. Immunophenotyping revealed loss of exhausted T cells and higher abundance of anti-cancer effector T cell populations. This effect was accompanied by a decrease of immunosuppressive lipid-laden macrophages and an expansion of CD14CD33high macrophages associated with antigen presentation. Analyses of human brain metastases samples supports a role of the ATP-adenosine signaling axis in modulating tumor inflammation and identified expression of CD39 and adenosine deaminase as predictive markers for patient survival and/or immune infiltration. Our findings demonstrate that the adenosine axis represents a druggable pathway to achieve local immunomodulation and treatment response, opening a new therapeutic avenue for brain metastases patients.

## Introduction

Brain metastasis (BrM) is the most common intracranial tumor in adults affecting 10 – 20% of solid cancer patients^1^. Treatment options remain limited and mostly comprise surgical resection, radio-and chemotherapy. Even with multimodality treatment, patient prognosis with an estimated 2-year survival rate of less than 10% is dire^1^. The introduction of immunotherapy in clinical practice has opened new therapeutic options for brain metastasis patients^2^. However, clinical benefit is limited in particular in patients with neurological symptoms^3^. Paucity in treatment success can be attributed to cancer cell intrinsic as well as niche-derived effects such as the highly immunosuppressive environment within intracranial lesions^4, 5^. Recent studies have shed light onto the importance of immunometabolism as a regulator of immune cell differentiation, function or activation and microenvironmental cues that pose a major metabolic barrier to effective anti-cancer immunity were identified^6^. Consequently, molecular drivers that modulate the status of tumor inflammation represent viable targets to facilitate anti-cancer immunity or to sensitize resistant tumors towards immunotherapy. The purine-adenosine signaling axis has been identified as an important metabolic checkpoint with immune modulatory functions^7, 8^. Activation of purine receptors P2X or P2Y by ATP or ADP triggers pro-inflammatory responses. In contrast, activation of adenosine receptors (A1R, A2AR, A2BR and A3R) by adenosine results in immunosuppression^9–12^. The ratio of purines to adenosine is controlled by the ectonucleotidase CD39 (encoded by *Entpd1*) that converts ATP to ADP and AMP and CD73 (encoded by *Nt5e*) that metabolizes AMP to adenosine. Adenosine concentrations are also controlled by adenosine kinase (ADK), which converts adenosine to AMP intracellularly and adenosine deaminase (ADA), which degrades adenosine to inosine. In contrast to adenosine, inosine has recently been linked to increased T cell stemness and fitness thereby improving efficacy of immune checkpoint blockade^13–15^. Hence, the purine-adenosine axis comprises multiple druggable components to stimulate anti-cancer immunity. Indeed, targeting of this signaling axis has shown anti-cancer efficacy^16, 17^. Here we show that adenosine signaling blockade synergizes with radiotherapy in a breast-to-brain metastasis mouse model and leads to modulation of the cellular composition within the tumor immune microenvironment evidenced by loss of exhausted T cells and expansion of effector T cell populations. Cellular changes were reflected by enrichment of pathways indicative of induction of adaptive immunity concomitant with a reduction of myeloid-mediated immunosuppression predominantly through interactions with lipid-laden macrophages. In patient BrM, we identified an adenosine signaling component signature that correlates with poor prognosis and reduced immune infiltration. In particular, CD39 showed negative correlation with immune cell infiltration, whereas reverse effects were observed for ADA. Collectively, our data demonstrates that perturbation of adenosine-mediated immunosuppression in combination with radiotherapy licences lymphoid and myeloid cells for anti-cancer immunity. This provides evidence for rational combination therapies to simultaneously modulate the lymphoid and myeloid compartment to overcome niche-specific limitations to therapy response in BrM.

## Results

### Abundance of purine-adenosine signaling components in brain metastasis

We first determined the concentrations of ATP, ADP, AMP, adenosine and inosine (Fig. 1a) in tumor-free brains and small and large tumor lesions of murine 99LN breast-to-brain metastases (99LN-BrM) using high performance liquid chromatography coupled to mass spectrometry (HPLC-MS/MS). In tumor-free brain, levels of adenosine exceeded the concentration of purines and inosine. Highest levels of ATP, ADP, AMP were detectable in brains harboring large tumors. Likewise, inosine levels were higher in tumor-bearing brains, whereas adenosine levels decreased compared to tumor-free brain. (Fig. 1b). To gain more detailed insight into the spatial distribution of ATP and adenosine in brain tumor and adjacent brain parenchyma, we performed mass spectroscopy imaging. This analysis revealed heterogenous distribution of ATP and adenosine in different brain areas and confirmed increased ATP levels and decreased adenosine levels in tumor lesions compared to adjacent brain (Fig. 1c,d). Next, we queried our previously generated RNA sequencing data (GSE164049) that contains bulk RNA sequencing data from fluorescence activated cell sorting (FACS)-purified immune cell populations harvested from 99LN-BrM as well as normal brain and blood from tumor-free mice^18^ to evaluate gene expression of purine-adenosine signaling components in tumor-associated and normal cells from control (Ctrl) mice. For the majority of analyzed signaling components, we observed increased gene expression in immune cells isolated from BrM (Fig. 1e), in particular in CD4+ and CD8+ T cells. Individual components showed decreased levels in tumor-associated (TA) immune populations such as *Entpd1* and *A3r* in TA-microglia (MG), *A2ar* in TA-Monocytes and TA-monocyte-derived macrophages (MDM) as well as the non-canonical adenosine pathway members *Enpp1* and *Entpd5*^19^ in TA-B cells. Analysis of BrM-associated immune cells in humans and other models of experimental BrM^20–24^ showed similar trends (Extended Data Fig. 1a,b). Single cell RNA sequencing (scRNAseq) of CD45+ immune cells from 99LN-BrM confirmed high *Entpd1*/CD39 expression in myeloid populations, whereas *Nt5e*/CD73 expression was largely confined to T lymphocytes (Fig. 1f and Extended Data Fig. 1c-e). Flow cytometric analysis corroborated the cell type-enriched expression pattern on protein level and detected high levels of CD73 and low levels of CD39 on 99LN tumor cells (Figure 1g). Analysis of a published spatial transcriptomic data set of human BrM (GSE179572)^23^ revealed expression of *ENTPD1*/CD39, *NT5E*/CD73 and *A2AR* in particular in immune cell rich areas (Extended Data Fig. S1f). Metabolomic analysis of freshly isolated and snap frozen patient glioma and BrM derived from breast, renal and lung cancer revealed higher levels of adenosine and inosine compared to ATP, ADP and AMP (Fig. 1h). In contrast, BrM derived from a melanoma patient showed higher ADP and AMP levels compared to adenosine. In summary, analysis of adenosine signaling components indicates a cell type-specific role in tumor-associated inflammation in mouse and human BrM.

**Figure 1.**
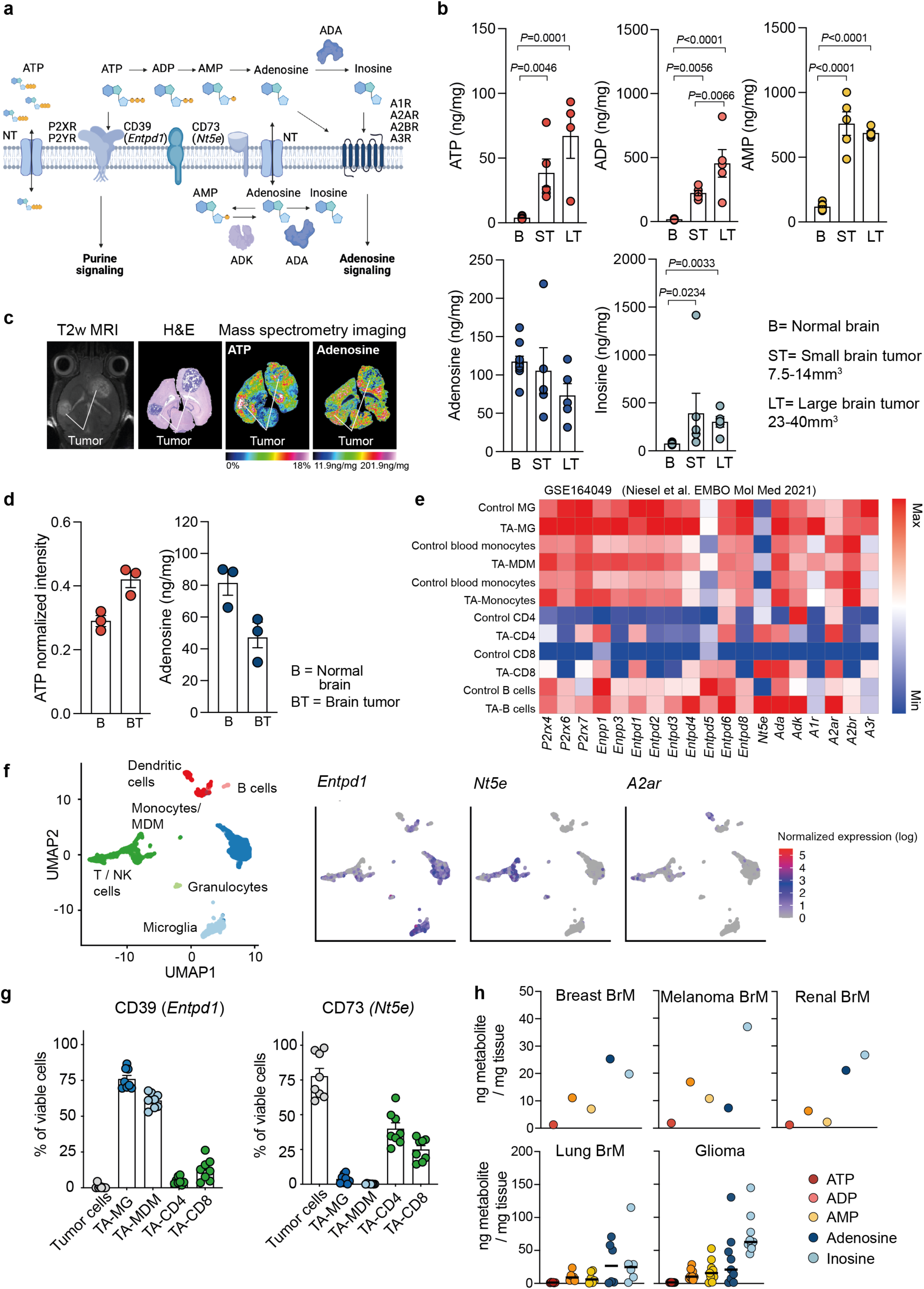
ATP-adenosine signaling axis in brain metastasis. **a,** Schematic overview of the ATP-adenosine signaling axis. **b,** Quantification of ATP, ADP, AMP, ADO and inosine (ng metabolite/mg tissue) in normal brain (n=10) and small (n=5) and large (n=5) 99LN-BrM based on liquid chromatography-mass spectrometry. Ordinary one-Way ANOVA with Tukey’s multiple comparisons test was used for ATP and ADP. Kruskal-Wallis with Dunn’s multiple comparisons test was used for AMP, adenosine and inosine. **c,** Representative images of mass spectrometry imaging of ATP and adenosine levels in the 99LN-BrM model. **d,** Quantification of mass spectrometry imaging of ATP and adenosine levels in normal brain and 99LN-BrM (n=3). Mann-Whitney log-rank was used. **e,** Heatmap depicts gene expression levels of signaling axis components in tumor-associated and control cells in the 99LN-BrM model. Data were obtained from Niesel *et al.* 2021 (Ref)^18^. **f,** Uniform manifold approximation and projection (UMAP) of scRNAseq data from sorted CD45+ cells showing *Entpd1*, *Nt5e* and *Adora2a* expression in the 99LN-BrM model. **g,** Frequency of CD39 and CD73-expressing cells were measured by flow cytometry in 99LN-BrM bearing mice (n=8). **h,** Quantification of ATP, ADP, AMP, adenosine and inosine in biopsies from glioma (n=7) and brain metastasis patients (n=1 for breast cancer, melanoma and renal; n=5 for lung).

### Genetic silencing of tumor-derived CD73 improves response to radiotherapy

Given the high levels of *Nt5e*/CD73 in 99LN tumor cells (Fig. 1g), we generated *Nt5e*-deficient 99LN cells using CRISPR/Cas9 (Extended Data Fig. 2a,b) and verified knockout efficiency on gene expression and enzymatic activity (Extended Data Fig. 2c-e). *Nt5e* deficiency in tumor cells did neither affect cell viability *in vitro* (Extended Data Fig. 2f) nor tumor incidence, progression and overall survival *in vivo* when injected into wildtype mice (Fig. 2a,b). Given previous reports on improved radiation response upon *Nt5e*/CD73 targeting^25^, we treated wildtype mice harboring *Nt5e* proficient and deficient 99LN-BrM with fractionated whole brain radiotherapy (WBRT) with 2Gy on 5 consecutive days (5 x 2Gy) using the small animal radiation research platform (SARRP)^26, 27^. Response to fractionated WBRT was more pronounced in animals with *Nt5e* deficient tumor cells leading to improved median survival rates (*Nt5e* KO 99LN WBRT) (Fig. 2b). To evaluate whether *Nt5e* deficiency in tumor cells increases response rates to immune checkpoint blockade, we treated mice with *Nt5e* deficient tumors with radiotherapy and/or neutralizing PD1 antibodies. Addition of immune checkpoint blockade by neutralizing PD1 antibodies did not significantly improve median survival rates (Fig. 2b) despite an increase in T cells and macrophage abundance (Extended Data Fig. 2g).

**Figure 2.**
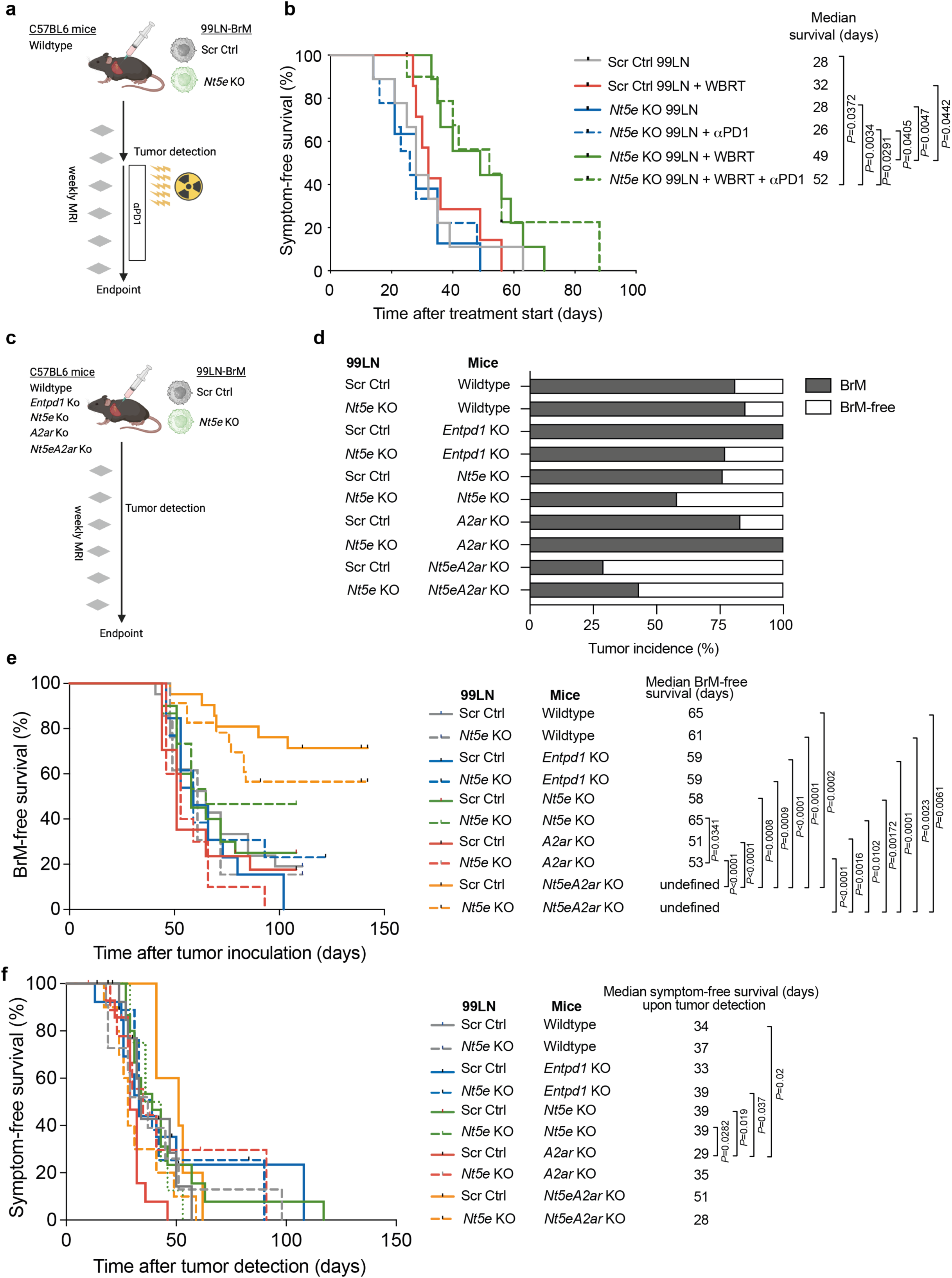
Targeting of *Entpd1/*CD3*9*, *Nt5e/*CD73 and *A2ar/*A2AR in combination with radiotherapy in murine brain metastasis. **a,** Scheme of the approach used to genetically target *Nt5e* in the tumor compartment in combination with whole brain radiation therapy (WBRT) in murine brain metastasis. **b,** Kaplan-Meier curves depict median of symptom-free survival (%) of the different groups: Scr Ctrl (28 days; n=9), Scr Ctrl + WBRT (32 days; n=8), *Nt5e*KO (28 days; n=9), *Nt5e*KO+*α*PD1 (26 days; n=9), *Nt5e*KO+WBRT (49 days; n=9), *Nt5e*KO+WBRT+*α*PD1 (52 days; n=9). Log-rank Mantel-Cox test was used. **c,** Scheme of the approach used to dissect the role of tumor cell-derived and stromal factors. **d,** 99LN-BrM incidence in the interrogated mouse strains: Scr Ctrl in Wildtype (n=21), *Nt5e*KO in Wildtype (n=13), Scr Ctrl in *Entpd1*KO (n=13), *Nt5e*KO in *Entpd1*KO (n=13), Scr Ctrl in *Nt5e*KO (n=20), *Nt5e*KO in *Nt5e*KO (n=15), Scr Ctrl in *A2ar*KO (n=17), *Nt5e*KO in *A2ar*KO (n=10), Scr Ctrl in *Nt5eA2ar*KO (n=21), *Nt5e*KO in *Nt5eA2ar*KO (n=23). **e,** Kaplan-Meier curves depict median of BrM-free survival (%) of the different groups: Scr Ctrl in Wildtype (65 days; n=21), *Nt5e*KO in Wildtype (61 days; n=13), Scr Ctrl in *Entpd1*KO (59 days; n=13), *Nt5e*KO in *Entpd1*KO (59 days; n=13), Scr Ctrl in *Nt5e*KO (58 days; n=20), *Nt5e*KO in *Nt5e*KO (65 days, n=15), Scr Ctrl in *A2ar*KO (51 days, n=17), *Nt5e*KO in *A2ar*KO (53 days, n=10), Scr Ctrl in *Nt5eA2ar*KO (undef, n=21), *Nt5e*KO in *Nt5eA2ar*KO (undef, n=23). Log-rank Mantel-Cox test was used. **f,** Kaplan-Meier curves depicted median of symptom-free survival (%) of the different groups: Scr Ctrl in Wildtype (34 days; n=16), *Nt5e*KO in Wildtype (37 days; n=11), Scr Ctrl in *Entpd1*KO (33 days; n=13), *Nt5e*KO in *Entpd1*KO (39 days; n=10), Scr Ctrl in *Nt5e*KO (39 days; n=15), *Nt5e*KO in *Nt5e*KO (39.5 days, n=8), Scr Ctrl in *A2ar*KO (29 days, n=14), *Nt5e*KO in *A2ar*KO (35 days, n=10), Scr Ctrl in *Nt5eA2ar*KO (51 days, n=5), *Nt5e*KO in *Nt5eA2ar*KO (28 days, n=10). Log-rank Mantel-Cox test was used.

### Genetic silencing of multiple adenosine signaling components reduces brain metastases incidence in the absence of radiotherapy

To distinguish between tumor-and host-derived effects, we injected *Nt5e*-proficient and deficient 99LN cells into wildtype, *Entpd1*KO, *Nt5e*KO, *A2ar*KO and *Nt5eA2ar*KO mice after confirming effects on purine and adenosine levels in tumor-free brains upon genetic silencing (Fig. 2c and Extended Data Fig. 2h). Genetic disruption of individual components in the tumor and/or host cells did not affect tumor incidence or progression (Fig. 2d-e). Only the injection of *Nt5e* KO tumor cells into *Nt5e* deficient mice resulted in reduced tumor incidence and *Nt5eA2ar*KO mice developed fewer 99LN-BrM, suggesting that multiple components or tumor-and host-derived sources should be targeted. Focusing on mice that developed brain metastases in the different experimental groups, we did not detect differences in tumor progression and median symptom-free survival time upon detection of established metastases (Fig. 2f). Quantification of immune cell infiltration showed no significant differences in the amount of CD3+, CD4+, CD8+ and IBA1+ cells in tumor-bearing animals across the analyzed conditions at trial endpoint when all mice had developed symptoms form brain metastases or the cumulative tumor size exceeded 100mm^3^ (Extended Data Fig. 2i).

### Pharmacological targeting of CD39, CD73 and A2AR synergizes with radiotherapy

To test the efficacy of pharmacological inhibition of adenosine signaling, we first tested the effect of single and combined inhibition of CD39, CD73 and A2AR compared to isotype treatment to choose the most efficacious regimen to block adenosine signaling in 99LN-BrM. We used the CD39 inhibitor POM1 (5 mg/kg), a CD73 neutralizing antibody (clone TY/23, 200 µg/mouse) and the A2AR inhibitor SCH58261 (10 mg/kg). Single CD39, CD73 or A2AR inhibitory treatment as well as combinations for double treatment (CD39iCD73i, CD39iA2ARi, CD73iA2ARi) did not result in significantly improved median survival (Fig. 3a). Only combined treatment with CD39iCD73iA2ARi resulted in significantly improved median survival compared to the isotype control group (Fig. 3a). We therefore chose to use combined pharmacological inhibition of CD39, CD73 and A2AR in the *in vivo* trials. We next tested the efficacy of this combination treatment with and without additional application of radiotherapy in 99LN-BrM. Using an independent cohort of mice, we confirmed that blockade of the adenosine signaling axis by combined inhibition of CD39, CD73 and A2AR leads to significantly improved survival compared to isotype-treated animals. Treatment of mice with fractionated WBRT in combination with adenosine signaling axis blockade resulted in synergistic effects leading to improved tumor control and significantly prolonged survival (Fig. 3b-d). Treatment combination with neutralizing PD1 antibody did not further improve median survival (Fig. 3d). despite a trend towards increased CD4+ and CD8+ T cell populations at trial end points (Extended Data Fig. 2j).

**Figure 3.**
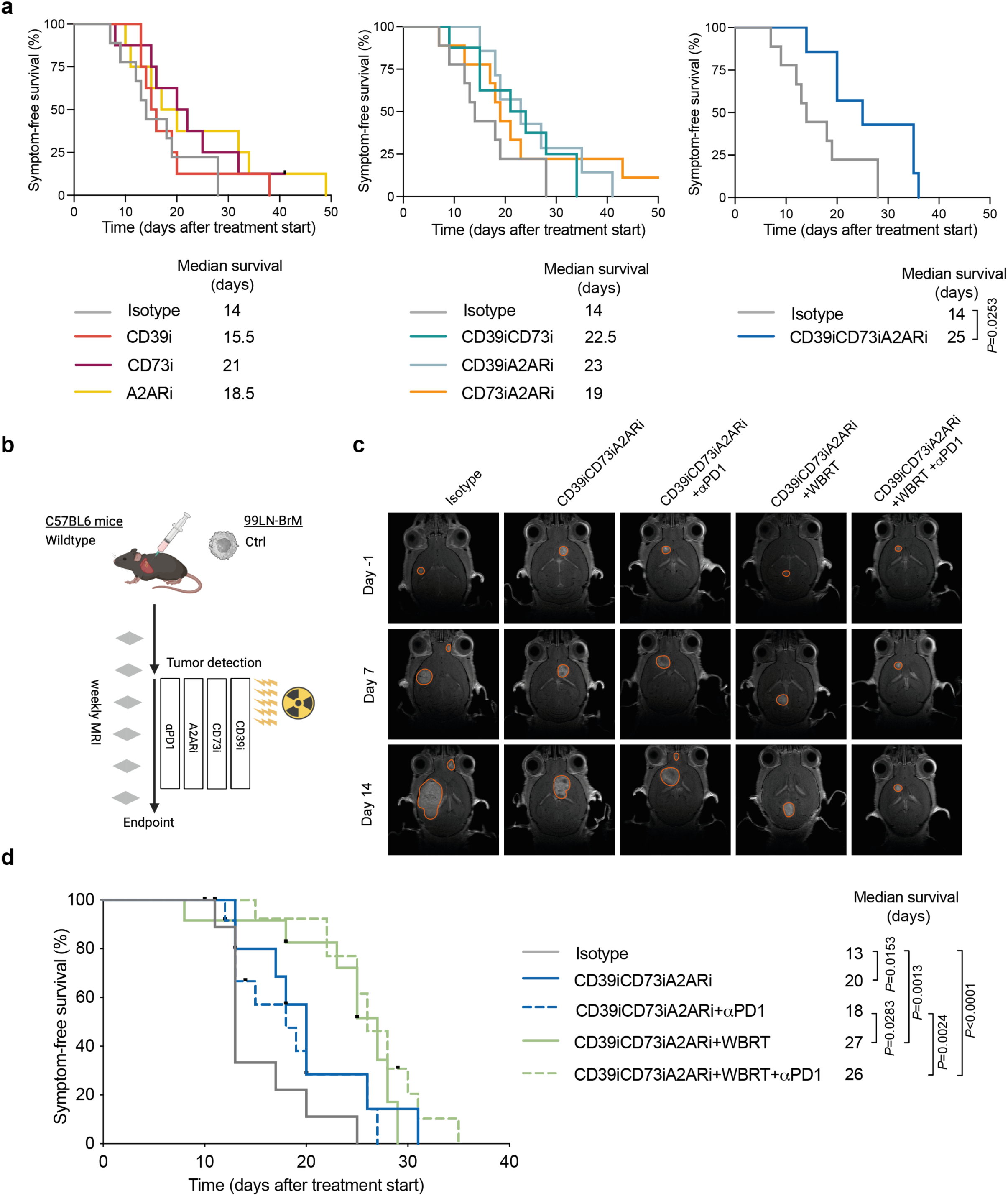
Pharmacological targeting of CD39, CD73 and A2AR synergizes with WBRT in murine breast-to-brain metastasis. **a,** Kaplan-Meier curves depict median of symptom-free survival (%) of the different groups: Isotype (14 days; n=9), CD39i (15.5 days; n=8), CD73i (21 days; n=8), A2ARi (18.5 days; n=8), CD39iCD73i (22.5 days; n=8). CD39iA2Ari (23 days; n=7), CD73iA2ARi (19 days; n=9) and CD39iCD73iA2ARi (25 days; n=7). Log-rank Mantel-Cox test was used **b,** Scheme of the pharmacological intervention trials and application of WBRT in experimental 99LN-BrM. **c,** Kaplan-Meier curves depict median of symptom-free survival (%) of the different groups: Isotype (13 days; n=9), CD39iCD73iA2ARi (20 days; n=13), CD39iCD73iA2ARi+*α*PD1 (18 days; n=13), CD39iCD73iA2ARi+WBRT (27 days; n=12), CD39iCD73iA2ARi+WBRT+*α*PD1 (26 days; n=13). Log-rank Mantel-Cox test was used. **d,** Representative MRI pictures (T2-weighted) of mice at day -1, 7 and 14 after treatment start belonging to the above-mentioned treatments.

### Adenosine signaling axis blockade combined with radiotherapy facilitates adaptive immunity

Even though we did not detect a correlation between immune cell abundance and treatment efficacy at trial endpoints, we sought to gain more detailed insight into the immune modulatory effects of adenosine signaling axis blockade with and without additional radiotherapy on the transcriptional level in different immune cell populations. We purified CD4+ and CD8+ T cells, B cells, MG, MDM and granulocytes as well as tumor cells using FACSorting and performed bulk RNA sequencing at day 15 after treatment start (Fig. 4a and Extended Data Fig. 3a). Evaluation of differentially expressed genes (DEG) indicated treatment-specific differences in transcriptional programs in tumor-associated immune populations and only minor changes in tumor cells (Extended Data Fig. 3b). We observed enrichment of immune modulatory genes including a range of cytokines, in particular in CD4+ T cells in isotype-treated mice compared to CD39iCD73iA2ARi+WBRT (*Ccl2, Ccl3, Ccl4, Ccl7, Ccl12, Cxcl9, Cxcl13, Cxcl16* and *Il1a*)*. Ccl7* and *Cxcl13* or only *Cxcl13* were enriched in CD39iCD73iA2ARi+WBRT compared to CD39iCD73iA2ARi or WBRT, respectively (Fig. 4b). Analysis of the cytokine abundance on tumor lysates containing tumor cells and tumor-infiltrating immune cells showed elevated levels of the majority of analyzed cytokines in the CD39iCD73iA2ARi and CD39iCD73iA2ARi+WBRT compared to isotype or WBRT (Fig. 4c-d). Levels of CXCL10, CXCL12 and CCL20 were increased in the CD39iCD73iA2ARi+WBRT group, whereas CXCL13, CCL4, CCL5 and IL16 were increased in CD39iCD73iA2ARi and CD39iCD73iA2ARi+WBRT animals. WBRT alone resulted in decreased levels of several cytokines compared to the other experimental groups, with only few cytokines that were enriched in response to WBRT (CX3CL1, CCL24, CCL27, IL2 and CCL17). CXCL1, associated with neutrophil and regulatory T cell recruitment^28^, showed increased abundance in response to isotype control compared to the other treatments. In concordance with the enrichment of cytokines associated with leukocyte recruitment^29^, we observed a significant increase of CD8+ T cells in CD39iCD73iA2ARi+WBRT treated animals compared to the other groups at the defined time point of 15 days after treatment start using flow cytometry (Fig. 4e-f). Functional annotation of DEGs revealed an induction of adaptive immunity in CD39iCD73iA2ARi+WBRT treated animals which was particularly apparent in the CD4+ T cell population with 19 out of 20 top pathways indicating immune cell recruitment and activation as well as antigen processing and presentation (Fig. 4g). CD8+ T cells, granulocytes and MDM showed similar enrichment of immune-related pathways in response to CD39iCD73iA2ARi+WBRT treatment (Extended Data Fig. 3c-e). CD8+ T cells showed additional alterations in L-glutamate import (Extended Data Fig. 3c) and MDMs showed additional alterations in pathways associated with metabolic processes including ATP metabolic processes and oxidative phosphorylation (Extended Data Fig. 3e).

**Figure 4.**
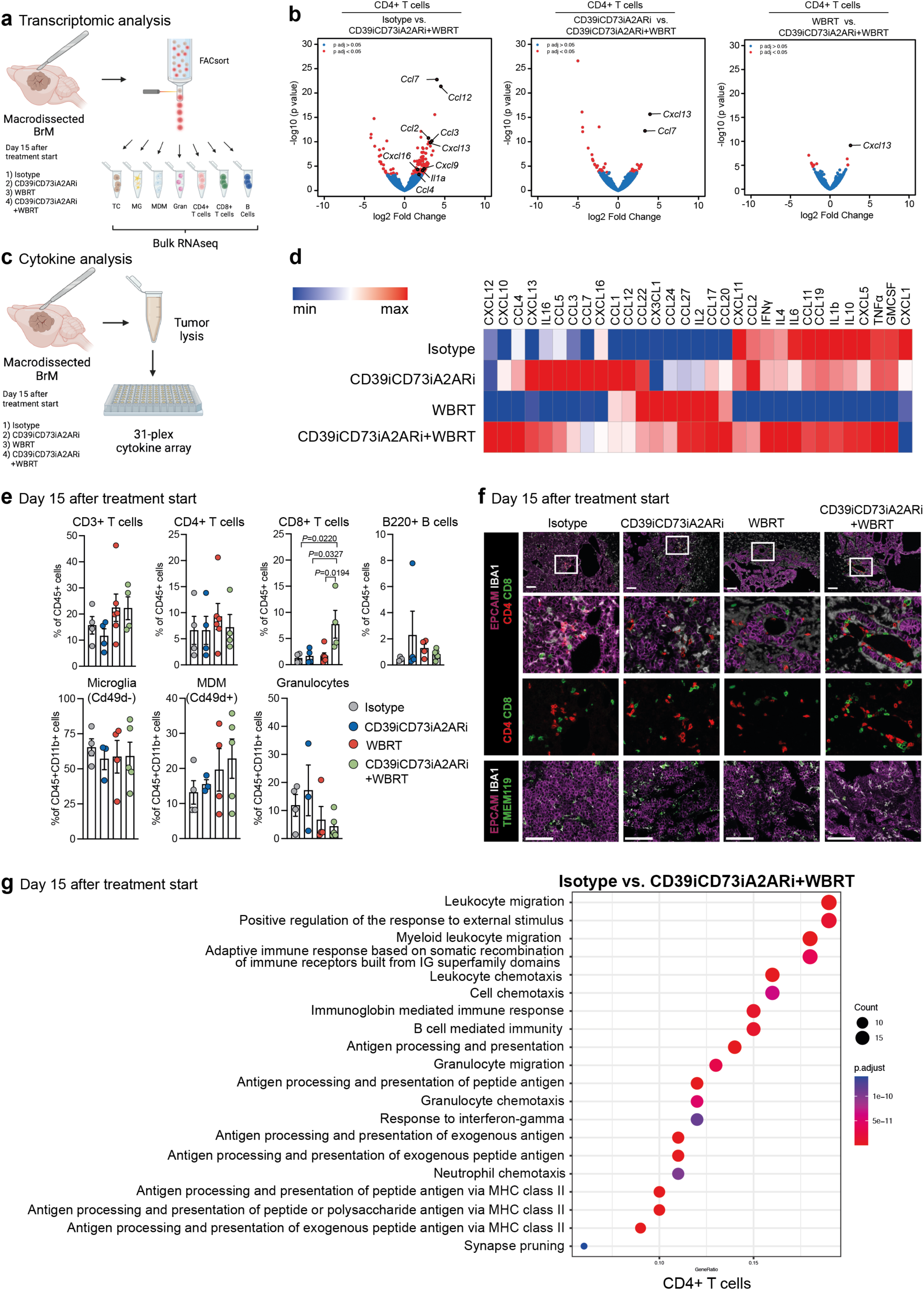
Immune modulatory effects of CD39iCD73iA2ARi+WBRT in established murine breast-to-brain metastasis. **a,** Schematic overview of FACS purified cell populations subjected to bulk RNAseq. **b,** Volcano plot depicts DEG highlighting the chemokines on sorted CD4+ T cells in the indicated comparisons. **c,** Overview of the approach to analyze the cytokine profiles in whole BrM lysates. **d,** Heatmap depicting the relative maximum and minimum values (arbitrary units) of cytokine concentrations in whole brain tumor lysates from 99LN-BrM mice treated with isotype (n=9), CD39iCD73iA2ARi (n=7), WBRT (n=9) and CD39iCD73iA2ARi+WBRT (n=10). **e,** Relative abundance of CD3+, CD4+, CD8+ T cells and B220+ B cells, microglia, monocyte-derived macrophages (MDM) and granulocytes in: isotype (n=4/4), CD39iCD73iA2ARi (n=4/3), WBRT (n=5/4) and CD39iCD73iA2ARi+WBRT (n=4/5); (lymphoid/myeloid panel). Ordinary one-Way ANOVA with Tukey’s multiple comparisons test was used. **f,** Representative images of 99LN-BrM stained for EPCAM (tumor cells), CD4 (CD4+ T cells), CD8 (CD8+ T cells), IBA1 (MDM/microglia), TMEM119 (microglia), Scale bar: 100 µm. **g,** Functional annotation of DEG (cut off padj < 0.05) on sorted CD4+ T cells from isotype-treated vs. CD39iCD73iA2ARi+WBRT-treated samples. Color gradient of red to purple indicates increasing padj; circle size indicates number of genes in pathway.

### Spatial heterogeneity of immune system-associated pathways in tumor core and stromal interface upon adenosine signaling axis blockade and radiotherapy

To gain insight into the cellular identity and functional association within different niches such as the tumor core and the tumor-stroma interface, we performed spatial transcriptomics with the GeoMx whole transcriptome atlas^30^ and used a segmentation method to distinguish PANCK+ tumor and PANCK-stromal regions. We stained formalin fixed and paraffin-embedded tissue sections with SYTO83, PANCK, IBA1 and CD3 to visualize nuclei, tumor cells, MDM/microglia and T cells. Histological assessment indicated a predominant localization of CD3+ T cells and IBA1+ MDM/microglia within stromal areas of tumor lesions in isotype, CD39iCD73iA2ARi and WBRT treated tumors. In contrast, CD39iCD73iA2ARi+WBRT treated tumors contained higher amounts of tumor cell-rich areas with direct interactions between CD3+ T cells and IBA1+ MDM/microglia (Fig. 5a,b). CD39iCD73iA2ARi+WBRT treated tumors contained a higher proportion of mature T cell populations, including CD8+ T effector cells (CD8T), gamma delta (γδ) T cells, NK T and NK cells and DC. The remaining treatment groups contained several populations of immature T cell progenitors denoted as double negative T cells (DN3 and DN4) and immature single positive cells (ISP) in the Imgen dataset^31^ used for deconvolution. CD39iCD73iA2ARi treatment resulted in accumulation of effector lymphocyte populations, in particular in the PANCK-stromal compartment (Fig. 5c). Changes in the proportions of myeloid cells, blood and lymphoid endothelial cells, pericytes and fibroblasts/astrocytes were less pronounced. Pathway annotation revealed enrichment of pathways indicating an activation of innate and adaptive immune responses in CD39iCD73iA2ARi+WBRT compared to isotype, CD39iCD73iA2ARi and WBRT (Fig. 5d-g and Extended Data Fig. 4a-e). While we observed enrichment of immune-related pathways in PANCK+ and PANCK-areas in CD39iCD73iA2ARi+WBRT compared to isotype and WBRT (Fig. 5d-g, Extended Data Fig. 4c,e), this effect was only apparent in PANCK+ areas in CD39iCD73iA2ARi+WBRT compared to CD39iCD73iA2ARi (Extended Data Fig. 4b,d). Pathways enriched in isotype and WBRT compared to CD39iCD73iA2ARi+WBRT were related to mitosis, cell cycle, chromosome maintenance and extracellular matrix remodeling (Fig. 5d-g, Extended Data Fig. 4c,e).

**Figure 5.**
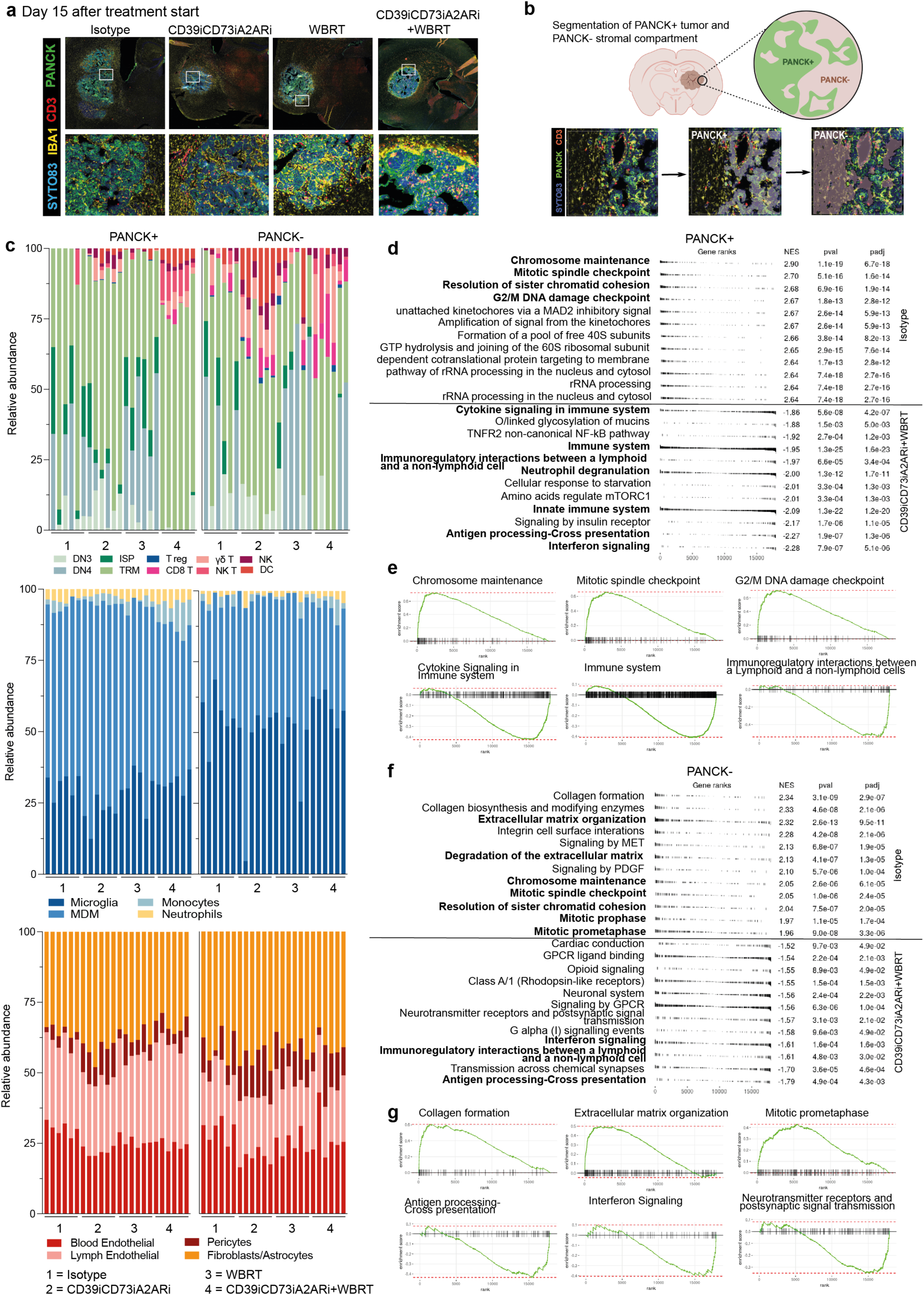
Spatial heterogeneity in immune modulation upon CD39iCD73iA2ARi+WBRT. **a,** Representative images showing PANCK (tumor cells), CD3 (T cells), IBA1 (MDM/microglia) and SYTO83 (nuclei staining) staining in the indicated treatment groups. **b,** Scheme depicting the approach employed for segmentation of tumor (PANCK+) and stromal (PANCK-) regions subjected to spatiaI transcriptomics. **c,** Cell abundances for the different conditions were estimated using the SpatialDecon R library. Related ROIs are grouped together and indicated on the x axis. **d,** Twenty pathways with lowest padj from the Molecular Signatures Database (MSigDB) Gene Ontology (GO) collection identified by gene set ORA in the DEG set isotype-treated vs. CD39iCD73iA2ARi+WBRT for PANCK+ areas. **e,** Mountain plots of some of the most differentially expressed pathways in isotype-treated vs. CD39iCD73iA2ARi+WBRT for PANCK+ areas. **f,** Twenty pathways with lowest padj from the MSigDB GO collection identified by gene set ORA in the DEG set isotype-treated vs. CD39iCD73iA2ARi+WBRT for PANCK-areas. **g,** Mountain plots of some of the most differentially expressed pathways in isotype-treated vs. CD39iCD73iA2ARi+WBRT for PANCK-areas.

### Adenosine signaling axis blockade together with radiotherapy reduces exhausted T cells and prevents immunosuppressive compensatory mechanisms

To further characterize shifts in immune populations, we employed scRNAseq on FACS purified CD45+ immune cells isolated from macro-dissected brain metastatic lesions (Extended Data Fig. 5a). After quality control, we identified 12,353 CD45+ cells, which were separated into T/NK cells, B and dendritic cells (DC), microglia and monocytes/MDM compartments based on marker gene expression (Fig. 1f and Extended Data Fig. 1c). Analyses of the relative proportion of each cell type within the CD45+ cell population revealed that in isotype and CD39iCD73iA2ARi tumors the predominant population were T and NK cells, followed by monocytes/MDM and minor contribution of microglia, granulocytes, B cells and dendritic cells. In contrast, the predominant cell population upon WBRT and CD39iCD73iA2ARi+WBRT were monocyte/MDM followed by microglia and T/NK cells. Proportion of granulocytes and B cells were similar across the experimental groups, whereas the proportion of dendritic cells increased in WBRT treated mice (Fig 6a). Differences in the relative proportions of cell populations could be explained by higher abundance of monocytes/MDM and microglia within the macro-dissected tumor lesion after irradiation as well as sensitivity of T/NK cells to radiotherapy^32^. To analyze effects on subpopulations within the lymphoid and myeloid compartments, we performed dimension reduction and unsupervised clustering separately for the individual compartments. Isotype and CD39iCD73iA2ARi treated tumors comprised high abundance of exhausted T cells (Fig. 6b,c and Extended Data Fig. 5b). CD39iCD73iA2ARi treatment resulted in minor population shifts leading to a relative expansion of NK T cells and Gzmk+ T cells, while exhausted T cells were less abundant compared to isotype-treated tumors. Radiotherapy led to an almost complete loss of exhausted T cells in WBRT and CD39iCD73iA2ARi+WBRT treated tumors. CD39iCD73iA2ARi+WBRT treated tumors showed an expansion of tissue resident memory T cells (TRM), Gzmk+ T cells and γδ T cells, whereas WBRT resulted in a predominant expansion of the NK T cell population followed by Gzmk+ T and γδ T cell populations (Fig. 6b,c). CD39iCD73iA2ARi, WBRT and CD39iCD73iA2ARi+WBRT treated tumors showed a relative increase of migratory DC (mig DC) clusters (Fig. 6d,e and Extended Data Fig. 5c). The TA-MDM compartment was clustered into monocytes, maturing MDM, Cxcl9 high MDM, lipid laden MDM, CD14CD33 high MDM and proliferating MDM (Fig. 6f,g and Extended Data Fig. 5d). Cxcl9 high MDM represented the most abundant subpopulation in isotype and CD39iCD73iA2ARi treated tumors. WBRT increased monocyte influx and expansion of lipid-laden MDM that have been associated with strong immunosuppressive functions^18, 33–35^, while CD39iCD73iA2ARi+WBRT resulted in expansion of CD14CD33 high MDM. CD14CD33 high MDM showed low expression of typical markers indicative of immunosuppressive or pro-inflammatory MDM functions and higher expression of markers associated with antigen presentation including CD81, H2-Eb and CD74 (Extended Data Fig. 5d). In isotype and CD39iCD73iA2ARi treated tumors, more than 85% of the MG population was stratified into a population resembling disease-associated microglia (DAM)^36^ (Fig. 6h,i and Extended Data Fig. 5e). We observed an expansion of homeostatic MG as well as Jun-Fos microglia in response to WBRT and microglia enriched for genes associated with mitochondrial metabolism in CD39iCD73iA2ARi+WBRT treated tumors. In WBRT and CD39iCD73iA2ARi+WBRT, we additionally identified microglia with stronger activation of the complement system.

**Figure 6.**
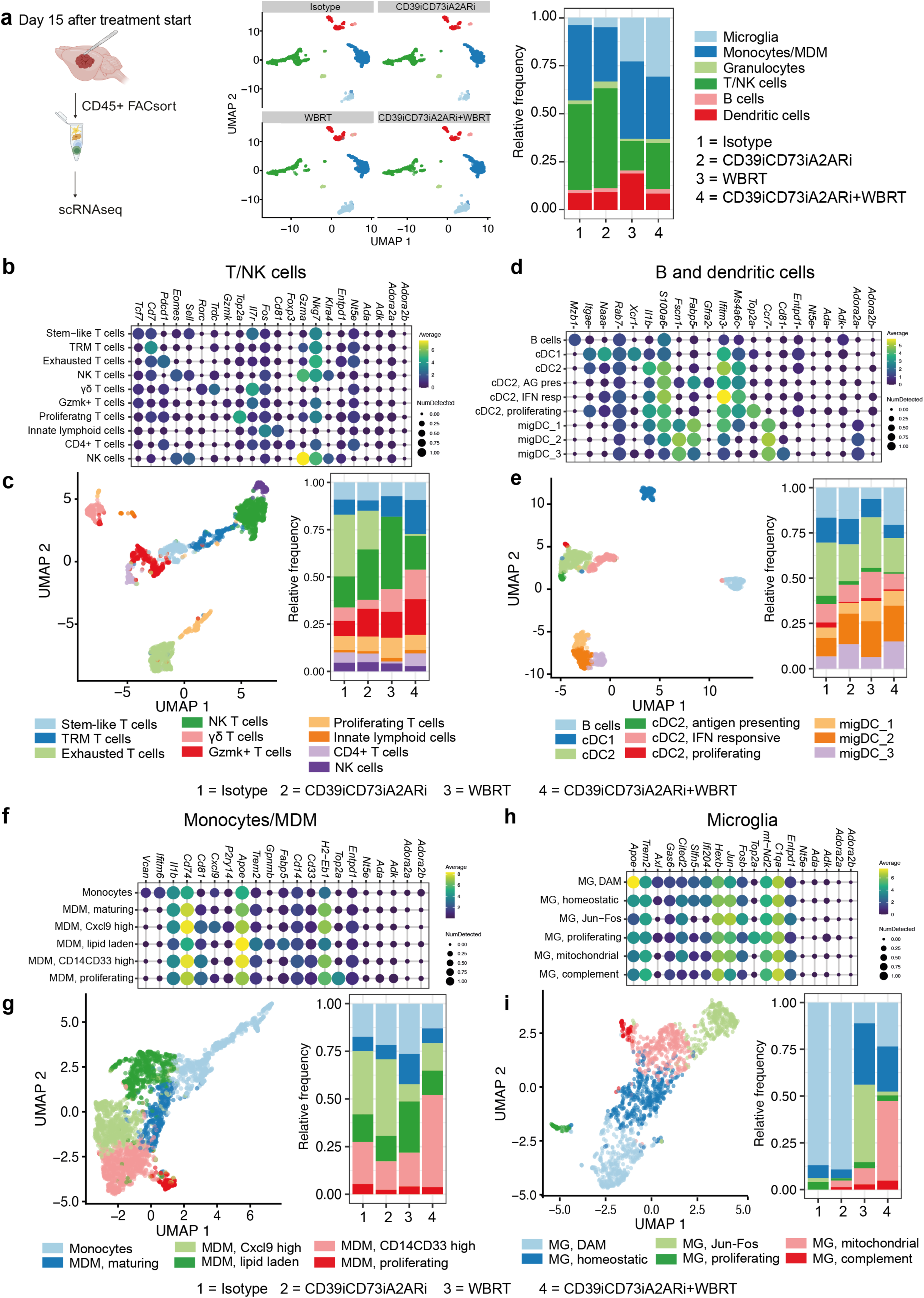
Population changes detected by scRNAseq upon CD39iCD73iA2ARi+WBRT in established murine 99LN-BrM. **a,** UMAP of scRNAseq from sorted CD45+ cells with cell compartment assignment and frequencies of cell compartment across the different treatment groups. **b,** Dot plot representing the discriminative marker genes for each functionally enriched state detected by scRNAseq for the T/NK cell compartment. Color indicates average expression value (logarithm of normalized counts). Size indicates the fraction of cells expressing the respective gene. **c,** UMAP of the T/NK cell compartment with subpopulation assignment and frequencies of subsets across the different treatment groups. **d,** Dot plot representing the discriminative marker genes for each functionally enriched state detected by scRNAseq for the B and dendritic cell compartment. **e,** UMAP of the B/dendritic cell compartment with subpopulation assignment and frequencies of subsets across the different treatment groups. **f,** Dot plot representing the discriminative marker genes for each functionally enriched state detected by scRNAseq for the macrophage/MDM compartment. **g,** UMAP of the monocyte/MDM compartment with subpopulation assignment and frequencies of subsets across the different treatment groups. **h,** Dot plot representing the discriminative marker genes for each functionally enriched state detected by scRNAseq for the microglia compartment. **i,** UMAP of the microglia compartment with subpopulation assignment and frequencies of subsets across the different treatment groups. 1 to 4 refers to 99LN-BrM mouse groups. 1=isotype, 2=CD39iCD73iA2ARi, 3=WBRT, 4=CD39iCD73iA2ARi+WBRT. Data obtained from CD45+ cells profiled by scRNAseq. n=3 mice per group were pooled before FACS purification.

### Analysis of cellular communication networks reveals changes in immunosuppressive versus stimulating interactions of myeloid cells and T cells

We next performed ligand receptor analysis using CellChat^37^ to dissect cell-cell interactions between T/NK cell populations and monocytes/MDMs across the different treatment conditions. This analysis revealed dominant interactions of monocytes/MDM populations with exhausted T cells in the isotype and CD39iCD73iA2ARi conditions (Fig. 7a). In WBRT treated animals, interactions were more evenly distributed across cell types with a particular shift towards interactions with NK T and NK cells likely due to the loss of exhausted T cells. In the CD39iCD73iA2ARi+WBRT condition, interactions were more frequently directed towards Gzmk+ T cells. Moreover, in WBRT treated tumors, lipid-laden macrophages showed dominant signal output compared to other MDM subpopulations (Fig. 7a). To further characterize cellular interactions, we queried ligand-receptor interactions between monocytes/MDMs and T cells focusing on cellular interactions that are involved in antigen presentation and T cell stimulation or suppression (Fig. 7b and Extended Fig. 6a). Monocytes, maturing MDM and in particular lipid-laden MDM showed highest probability of T cell suppressing interactions including interactions of *Spp1* and *Fn1* with *Cd44* and integrins (e.g. *Itgv3*, *Itgb1*, *Itgb3* and *Itga4*) as well as *Thbs1*-*Cd47* interactions^38–40^ (Fig. 7b and Extended Data Fig. 6c). Among those, lipid-laden macrophages showed highest probability of mediating T cell suppressive function via *Spp1*. Monocytes and lipid-laden macrophages in isotype and WBRT samples showed no probability of *Icos*-mediated modulation of T cell functionality, whereas the CD39iCD73iA2ARi and CD39iCD73iA2ARi+WBRT group retained *Icos*-mediated interactions in both subpopulations. With regard to antigen-presentation, *MHC class I-Cd8* and *MHC class II-Cd4* interactions occurred with highest probability towards exhausted T cells and CD4+ T cells followed by proliferative T cells and Gzmk+ T cells in isotype treated tumors (Fig. 7b and Extended Data Fig. 6c). Only minor changes were observed in CD39iCD73iA2ARi treated tumors. In CD39iCD73iA2ARi+WBRT tumors, interactions shifted towards Gzmk+ T cells (Fig. 7b and Extended Data Fig. 6c). In contrast, WBRT treatment alone resulted in decreased probability of interactions indicative of antigen-presentation mostly feeding into TRM and Gzmk+ T cells (Fig. 7b). Within the MG compartment, immunosuppressive interactions in particular by *Lgals9*-*Cd45* as well as interactions of *Icam1* with *Itgal* and *Spn* previously associated with leukocyte migration as well as positive and negative T cell activation^41, 42^ became more dominant in the WBRT and CD39iCD73iA2ARi+WBRT treatment groups (Fig. 7c and Extended Data Fig. 6b).

**Figure 7.**
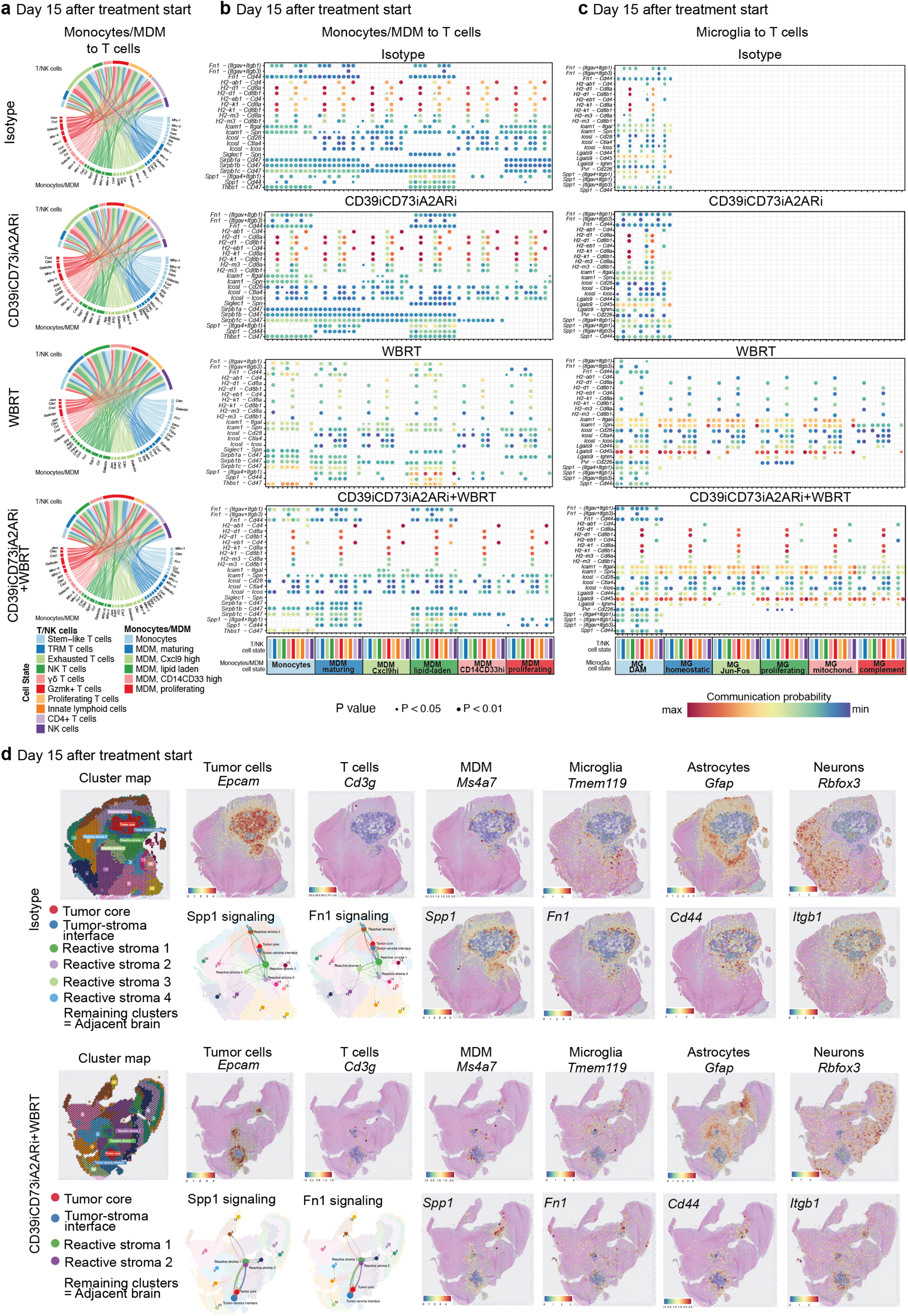
Cell-cell interaction analysis between the T/NK compartment and monocytes/MDM and microglia in 99LN-BrM. **a,** Chord diagram showing the cell-cell interaction of selected pathways. The perimeter of the chord diagram is color-coded by cellular subtype (colored outer arcs). The inner connecting lines show the interactions between target and receiver cell types. The connecting lines are color coded consistently with cellular subtypes according to the sending cell type. The thickness of each edge (edge weight) is proportional to the interaction strength. **b,** Bubble plot depicting in the y axis the ligand-receptor pair interactions and in the x axis the interacting T/NK and monocytes/MDM cell subclusters from the isotype condition. Color gradient of blue to red shows the communication probability. Increased circle size indicates decreased p value. Only interactions with a p-value < 0.05 are shown. **c,** Bubble plot depicting in the y axis the ligand-receptor pair interactions and in the x axis the interacting T/NK cell subclusters and microglia from isotype, CD39iCD73iA2ARi, WBRT and CD39iCD73iA2ARi+WBRT. Color gradient of blue to red shows the communication probability. Increased circle size indicates decreased p value. Only interactions with a p-value < 0.05 are shown. **d,** Representation of spatial transcriptomic spot clustered according to their transcriptomic profile. The expression of cell-type specific marker genes (as log2 normalized counts) is shown in superposition to an HE staining of the tissue section.

To further characterize immunosuppressive interactions of microglia or monocyte/MDM with T cells, we performed spatial transcriptomics using 10x Visium. Expression of *Spp1*, *Fn1* and its interaction partners *Cd44* and *Itgb1* was enriched within spatial transcriptomic clusters identifying the tumor-stroma interface and reactive stroma (Fig. 7d). A similar distribution was observed for *Thbs1* expression, while *Cd47* expression was detected across the brain parenchyma (Extended Data Fig. 6d). *Spp1* signaling was identified as one of the top enriched pathways and visualization of interaction probability showed dominant interactions within the tumor core, tumor-stroma interface and reactive stroma. Similar patterns were observed for *Fn1* and *Thbs1* signaling indicating an induction of immunosuppressive interactions directed towards tumor-associated immune cells that are not present in the adjacent brain parenchyma (Fig. 7d and Extended Data Fig. 6d). Therefore, the spatial transcriptomic data confirms the probability of receptor ligand interactions identified in the scRNAseq data given their co-expression within BrM lesions and reactive stroma. Taken together, multiomics immunophenotyping identified cellular and molecular changes within the immune microenvironment. In particular, adenosine signaling blockade in combination with radiotherapy resulted in immunomodulation that supports anti-cancer immunity but prevents myeloid-mediated immunosuppression triggered by WBRT as monotreatment.

### Abundance of adenosine signaling components associates with prognosis and correlates with immune infiltration in patient brain metastasis

To evaluate the translational significance of our experimental data, we determined the histological score based on expression levels and amounts of cells stained for CD39, CD73, A2AR, A2BR, ADA and ADK in tumor cells on tissue microarrays (TMA) containing 244 brain metastatic tumors from melanoma, non-small cell lung cancer (NSCLC), small cell lung cancer (SCLC) and breast, colon and renal cancer (Extended Data Fig. 7a,b). We stratified patients based on median split for each factor and found association of Cluster 1, defined by high CD39 and A2BR, with poor survival rates and overall lower immune score (Fig. 8a-d). Further evaluation revealed an association of high CD39 expression with high proliferation rates (Ki67+) and decreased MDM/microglia (IBA1+) and CD8+ T cells. In contrast, ADA showed a reverse correlation with proliferation rates and a positive correlation with CD8+ and MDM/microglia accumulation (Fig. 8e), whereas no significant correlations were observed for A2BR. Stratifying patients based on their CD39 and ADA expression revealed an association of CD39^high^ADA^low^ expression with poor survival rates compared to CD39^low^ADA^high^ patients (Fig. 8f). A similar distribution of BrM patients from renal, breast cancer and NSCLC were grouped in the CD39^high^ADA^low^, mixed (CD39^high^ADA^high^ and CD39^low^ADA^low^) and CD39^low^ADA^high^ groups, while a higher proportion of melanoma patients were in the CD39^low^ADA^high^ group. In contrast, patients with BrM derived from colon or SCLC were mostly classified in the CD39^high^ADA^low^ group (Fig. 8g). We observed that high CD39 expression alone correlates with poor prognosis (Fig. 8h). In contrast, ADA expression alone is not predictive for survival (Fig. 8k). The human data suggests a role of the adenosine axis in BrM as well as BrM-associated inflammation and indicates potential differences in the balance of purine-driven inflammation vs. adenosine-mediated immune suppression in BrM derived from different primary cancer entities.

**Figure 8.**
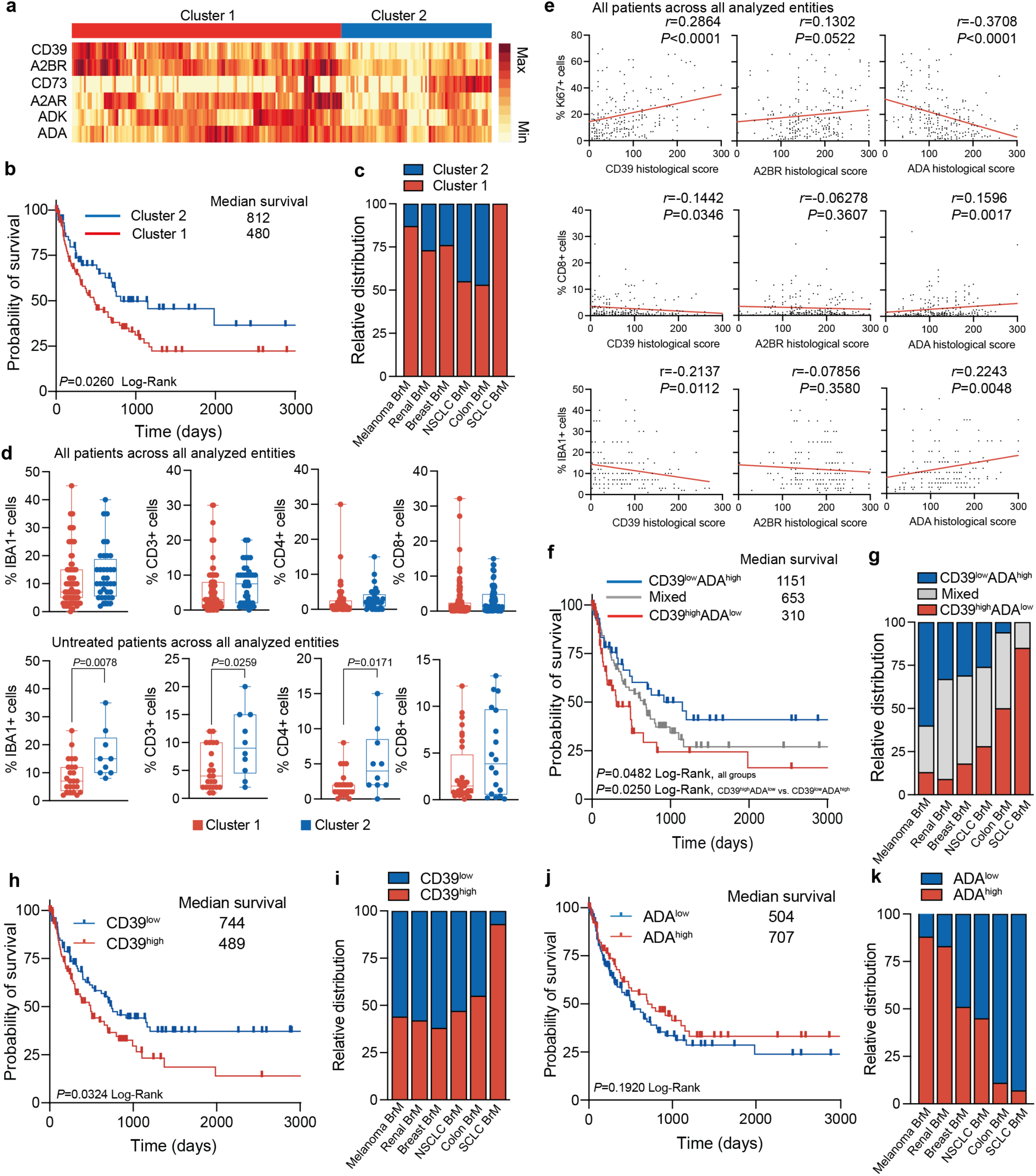
Assessment of ATP-adenosine components in human brain metastasis. **a,** Heatmap showing histological scores for CD39, A2BR, CD73, A2AR, ADA and ADK in patient brain metastasis across different tumor entities that drive the cluster aggrupation 1 and 2. Cluster 1 (n=128); cluster 2 (n=70). **b,** Kaplan-Meier curve shows probability of survival upon brain metastasis detection for patients in cluster 1 and 2. Log-rank Mantel-Cox test was used. **c,** Relative distribution into cluster 1 and 2 across brain metastasis patients from the different cancer types. **d,** Representation of the correlation with IBA1, CD3, CD4 and CD8 infiltration for patients belonging to cluster 1 and 2. Mann-Whitney t test was used. **e,** Correlation of tumor proliferation (Ki67) and CD8 and IBA1+ immune infiltration with CD39, A2BR and ADA histological score in tumor cells across brain metastasis samples from all tumor entities. Pearson correlation test was used. **f,** Kaplan-Meier curve shows probability of survival upon brain metastasis detection for patients showing a tumor signature composed of CD39^low^ADA^high^ vs. CD39^high^ADA^low^. Log-rank Mantel-Cox test was used. CD39^low^ADA^high^ (n=56); CD39^high^ADA^low^ (n=63) and mixed (n=102). **g,** Relative distribution into the different signatures across brain metastasis patients from the different cancer types. **h,** Kaplan-Meier curve shows probability of survival upon brain metastasis detection for patients showing high or low CD39 expression. Log-rank Mantel-Cox test was used. CD39^high^ (n=114) and CD39^low^ (n=120). **i,** Relative distribution of CD39^high^ and CD39^low^ tumors across brain metastasis patients from the different cancer types. **j,** Kaplan-Meier curve shows probability of survival upon brain metastasis detection for patients showing high or low ADA expression. Log-rank Mantel-Cox test was used. ADA^high^ (n=101) and ADA^low^ (n=124). **k,** Relative distribution of ADA^high^ and ADA^low^ tumors across brain metastasis patients from different cancer types.

## Discussion

Different myeloid-and lymphoid-targeted strategies have been tested in preclinical models to convert the immunosuppressive tumor microenvironment into a proinflammatory milieu and to reactivate exhausted T cells^18, 21, 35, 43^. However, compensatory mechanisms often blunt the efficacy of such strategies. This suggests that simultaneous modulation of disease-associated activation and exhaustion states in myeloid and lymphoid cells is key to facilitate prominent anti-cancer immunity and to overcome the induction of adaptive resistance to immune-targeted therapy. Our study exemplifies that targeting the adenosine signaling axis combined with radiotherapy affects the myeloid and lymphoid compartment and represents a viable strategy to promote anti-cancer immune responses in experimental BrM. The ATP-adenosine axis has been identified as an immunomodulatory pathway in previous studies. In extracranial tumors, combined targeting of convertases and adenosine receptors improved tumor control and enhanced T cell infiltration into tumors compared to individual adenosine pathway component targeting^17^. Our data confirms the benefit of combined targeting of purine convertases and adenosine receptors. Yet, the most prominent effects are observed upon targeting of the adenosine signaling axis in combination with radiotherapy applied as fractionated image-guided WBRT in 99LN-BrM. Synergistic effects could be attributed to (1) a potential link of CD73 expression in tumor cells to radio-resistance^25^, (2) the necessity to increase extracellular ATP by ionizing radiation to shift the signaling cascade towards purine-driven inflammation^44^, (3) the clearance of exhausted T cell populations for reconstitution of the T cell niche with anti-cancer effector cells and (4) the simultaneous modulation of the lymphoid and myeloid compartment to prevent myeloid-mediated immunosuppression^45^. In this study, we focused on dissecting immunomodulatory effects of radiotherapy in combination with adenosine signaling axis blockade. Using scRNAseq, we detected similar abundance of T cells with a stem-like signature across the different preclinical treatment conditions and loss of exhausted T cells upon irradiation, in concordance with a recent study in human BrM from a NSCLC patient^46^. Stem-like T cells are reported to be less radio-responsive^47^ and this may explain their consistent proportion across the different conditions. In CD39iCD73iA2ARi+WBRT-treated animals, the T cell niche shifted towards higher presence of populations associated with prominent anti-cancer activity^48–51^ such as tissue resident memory T cells, γδ T cells and Gzmk+ effector T cells, whereas WBRT alone led to an expansion of NK T and γδ T cells. Gene signatures suggest an enrichment of transcriptional programs indicative of CD4+ T cell interactions with antigen presenting cells and signatures associated with adaptive immune responses in stromal and cancer regions in CD39iCD73iA2AR+WBRT-treated animals. An association with antigen presentation was also evident in the monocyte/MDM population marked by an expansion of CD14CD33hi MDMs that showed increased expression of genes associated with antigen presentation.

While radiotherapy alone also resulted in pronounced changes within the T/NK cell compartment, the influx of immunosuppressive MDM counteracts the induction of T cell responses. Compensatory effects mediated by immunosuppressive MDM populations has previously been reported to diminish T cell activation and accounts for poor response rates to immune check point blockade in preclinical models and patient cohorts ^18, 33, 35^. In particular, lipid-laden macrophages have been identified as highly immunosuppressive ^33, 34^. Our data confirmed the expansion of lipid-laden macrophages in response to radiotherapy and identified receptor-ligand interactions that are involved in myeloid-mediated immunosuppression. In future studies, we seek to functionally evaluate the role of immunosuppressive receptor-ligand interactions between macrophages and T cells to specifically resolve lipid-laden macrophage-mediated T cell suppression. In particular *Spp1* represents an interesting target given its correlation with unfavourable outcomes^52, 53^.

The prognostic impact of CD39, CD73, A2AR and A2BR has previously been assessed in tumors such as breast cancer, melanoma and NSCLC^54–57^. However, only one study has reported CD39 and CD73 expression in melanoma brain metastasis, and similar to our findings no correlation was found between CD73 and patient survival^58^. In our study, histological evaluation of patient BrM revealed a correlation of CD39 expression in tumor cells with poor prognosis, increased tumor cell proliferation and reduced immune cell infiltration, whereas ADA expression showed inverse correlations with respect to tumor cell proliferation and positive correlation with immune infiltration. Even though CD39 expression could be a valuable prognostic marker for BrM patients, analysis of multiple markers, such as the signature comprising CD39, CD73, A2AR, A2BR, ADA and ADK showed more robust results and indicated a dependency on the primary tumor entity that metastasized to the brain. Moreover, our data revealed that combining protein expression levels of CD39 and ADA stratified patients based on clinical outcome, immune infiltration and association to specific brain metastatic primary tumor entities. In this regard, SCLC and colon brain metastasis patients clustered in the poor prognosis group with high CD39 and low ADA expression. This indicates that patients with strong adenosine signaling signatures might benefit more from pathway blockade than patients with signatures indicative of purine signaling. However, it also has to be mentioned that previous treatment history might affect the composition of the immune infiltrate and the expression levels of purine-adenosine signaling axis components, which needs to be considered to assess their applicability as predictive markers. Furthermore, we evaluated the histological score based on expression levels in tumor cells. Given the high expression of purine-adenosine signaling components in immune cells, follow-up analysis should interrogate correlations of the prognostic value of immune cell-specific expression of CD39, CD73, A2AR, A2BR, ADA and ADK. Indeed, CD39 is regarded as an indicator of tumor-reactive or tissue-resident CD8+ T cells^59–61^. The expression of CD39 in distinct T cell subsets in different cohorts of breast cancer correlates with better patient prognosis^62, 63^, although opposite findings are reported for metastatic melanoma treated with immunotherapy^64^.

Targeting the adenosine pathway may further potentiate immune responses induced by radiotherapy and immunotherapy^8^ e.g. by immune checkpoint blockade. However, in the 99LN-BrM model, we did not observe improved tumor control and survival upon PD1 inhibition in combination with adenosine pathway blockade, independent of WBRT. This may be due in part to the loss of the CD8+ PD1+ population (exhausted T cells) in response to WBRT as shown in this study. Current clinical trials with breast cancer patients combine CD73 blockade with chemotherapy, radiotherapy or immune checkpoint blockade (NCT03875573 and NCT03616886)^65, 66^. Important to note is the exclusion of patients with metastatic disease from those trials and that results from studies focussing on extracranial disease cannot be easily transferred to brain metastases.

Using preclinical models, we propose that adenosine signaling axis blockade needs to be combined concurrently with WBRT to improve therapeutic efficacy in BrM. Our study provides insight for the first time into purine-adenosine axis-mediated modulation of brain metastasis-associated inflammation and thereby opens new avenues for metabolic checkpoint blockade to simultaneously modulate tumor-infiltrating myeloid and lymphoid cells to facilitate anti-cancer immunity.

## Author contributions

AS-B, and LS conceptualized the study. AS-B, MS, JM, JA, AM, IT, DM, JH, JM, BR, YR, GH, DB, HM, MBR, MH and LS performed experiments, analyzed data and/or provided guidance on omics analysis or multiplexed histology. SH and FR assisted with the application of radiation therapy. MS, JM, JS, MHM, PNH, KI performed computational analysis. TB, MC, KHP, and PNH provided patient samples. AS-B and LS wrote the manuscript. All authors edited or commented on the manuscript. LS supervised the study.

## Acknowledgments

We thank all individuals who donated brain tumor tissue. We also thank Petra Dinse, Tijna Alekseeva, Stefan Stein, Annette Trzmiel, Michelle Antonios, Jeannie Peifer and Jennifer Lun for technical support. We also thank Matthias Ebert and animal caretakers at Georg-Speyer-Haus. We thank Verena Jendrossek for providing CD39-knockout mice and Johanna A. Joyce for providing the 99LN cancer cell line. AS-B was recipient of the EACR-NanoString-Illumina Whole Transcriptome Grant. K.I. received funding from the Mildred Scheel Career Center Frankfurt (German Cancer Aid). Research in the lab of LS is supported by institutional funds from the Georg-Speyer-Haus jointly funded by the German Federal Ministry of Health and the Ministry of Higher Education, Research and the Arts of the State of Hesse, as well as grants from the German Research Foundation (SE2234/3-1and SE2234/3-2), the German Cancer Aid (Max-Eder Junior Group Leader Program 70111752 and 70114437), the LOEWE Center Frankfurt Cancer Institute (FCI), the Excellence Cluster iFIT (EXC2180), the HECTOR Foundation MINT Personal Fonds, the German Cancer Consortium (DKTK, partner site Frankfurt/Mainz), the Beug Foundation for Metastasis Research and the Dr. Bodo Sponholz Foundation. Scientific illustrations were created using BioRender.com.

## Declaration of interests

The authors declare no competing interests.

## Data availability

Bulk RNA sequencing, single cell RNA sequencing and spatial transcriptomics (GeoMx and Visium) data have been deposited in GEO (superseries GSE248307). Metabolomics data have been deposited in the EMBL-EBI MetaboLights database (MTBLS8929)^67^. This paper analyzes existing, publicly available data and the accession numbers for these datasets is listed in the corresponding figure as well as in the online methods. Microscopy data reported in this paper will be shared by the lead contact upon request. Reagents generated in this study will be shared with other research groups only upon receipt of reasonable request on a case-by-case basis with a completed Materials Transfer Agreement (MTA). Any additional information or data reported in this manuscript will be available from the lead contact upon request.

## Online Methods

### Clinical samples

All participants included in this study provided written consent and did not receive compensation for their participation. Formalin-fixed and paraffin-embedded tissue from archived BrMs collected between 2003 and 2018 was processed as tissue microarrays (TMAs) and provided by P.N.H. and K.H.P. Specimens were obtained from the UCT tumor bank (Goethe University, member of the German Cancer Consortium and German Cancer Research Center). The TMAs included specimens from 244 patients. To reduce selection bias, patient recruitment was performed by clinical staff that was not directly involved into the design and analysis of this study. Approval for this study was conferred by the ethics committee UCT Frankfurt/Goethe University, project numbers GS4/09, SNO-01-12, SNO-02-2015 and SNO-05-2018. For metabolomic analyses, tissue from BrM and glioma was fresh frozen in liquid nitrogen and stored at -70°C until further processing. Approval was conferred by the ethics committee UCT Frankfurt/Goethe University, project number UCT-2-2022. Final inclusion of clinical samples was based on pathological diagnosis performed outside the Sevenich laboratory. Additional information about the clinical samples can be found in Supplementary Table 1 and ^68^.

### Animal studies

Animal studies were approved by the local government committees (Regierungspräsidium Darmstadt, Germany protocol number F123/1068) and experiments were conducted in accordance with all ethical requirements at the Georg-Speyer-Haus. Mice were maintained in a temperature-controlled room under standard dark diurnal cycles. They were housed in filter-topped cages (maximum 5 mice per cage) and fed standard laboratory chow and water *ad libitium*. C57BL6J (IMSR_RJ:C57BL-6JRJ), *Nt5e*KO (IMSR_JAX:018986) and *A2ar*KO (IMSR_JAX:010685) mice were obtained from Charles River. The *Entpd1*, the *Nt5e* and *A2ar*KO mouse lines were generated as described previously^69–71^. *A2ar*KO mice were crossed to the C57BL6J strain for 6 generations and were tested for genetic integrity using the Transnetyx Genetic Monitoring. The *Nt5eA2ar*KO mouse line was generated by cross-breeding and tested using the Genetic Monitoring platform from Transnetyx. The *Entpd1A2ar*KO, the *Entpd1Nt5e*KO and *Entpd1Nt5eA2ar*KO mice lines were generated by cross-breading. Animal genotyping was performed by Transnetyx using real-time PCR.

### Cell lines

The murine parental 99LN cell line was derived from a metastatic lymph node of the MMTV-PyMT breast cancer model (C57BL6J background) and twofold selected *in vivo* for brain homing capacity as previously described^72^, resulting in the 99LN variant used herein. The 99LN and HEK293T (RRID:CVCL_0063) cell lines were maintained in DMEM containing 10% fetal bovine serum with 1% L-glutamine and 1% penicillin/streptomycin.

### Generation of *Nt5e*-deficient tumor cells

The lentiviral vectors used to knockout *Nt5e* in our study, pLV[CRISPR]-hCas9:T2A:Bsd-U6>mNt5e[gRNA#2944]-U6>mNt5e[gRNA#3333] (VB180408-1067rnu) and pLV[CRISPR]-hCas9:T2A:Bsd-U6>mNt5e[gRNA#2934]-U6>mNt5e[gRNA#3335] (VB180410-1343wdp), were constructed and packaged by VectorBuilder. pLV[CRISPR]-hCas9:T2A:Bsd-U6>scrambled_gRNA1-U6>scrambled_gRNA2] (VB180416-1215auc) was used as a scrambled control. Supplementary Table 2 contains further information on the constructs used. Virus was generated following the manufacturer’s instructions of ProFection® Mammalian Transfection System Kit (Promega, Cat#E1200) using the HEK293T cells and the envelope and packaging vectors pMD2.VSVG (Addgene, RRID: Addgene_12259) and pSPAX2 (Addgene, RRID: Addgene_12260). Cells were selected for plasmid integration (blasticidin) and sorted based on CD73 expression using a CD73-specific antibody (CD73-VioBright 515, Miltenyi Biotec, Cat#130-111-335, REA778, RRID: AB_2659163). *In vitro* assays were performed with *Nt5e* KO1 and KO2 99LN cells. *Nt5e* KO2 99LN cells were used for *in vivo* experiments.

### Generation of experimental brain metastasis

For brain metastasis generation, 1 x 10^5^ 99LN cells were inoculated into the left cardiac ventricle of 10-12 week-old female C57BL6J and 8-14 week-old female and male *Entpd1*KO, *Nt5e*KO, *A2ar*KO and *Nt5eA2ar*KO mice. To induce intracranial tumors, 5 x 10^4^ 99LN cells were injected 2.5 mm to the left of bregma, 0.4 mm posterior and 3 mm below the surface of the skull using a stereotactic frame (Stoelting). All *in vivo* data for flow cytometry, FACS sorts, RNAseq and *in vivo* trials were generated from multiple biological replicates per group (see figure legends).

### *In vivo* MRI measurements

Tumor progression was monitored by magnetic resonance imaging (MRI) measurements. MRI was performed using a 7 Tesla Small Animal MR Scanner (PharmaScan, Bruker) with a volume coil as transmitter and head surface coil for signal reception. Mice were injected intraperitoneally with 150 µl gadobutrol (Gadovist, 1 mmolml-1, Bayer) before imaging. Data acquisition was performed using Paravision v.6.0.1 software with images acquired in coronal planes. For T2-weighted images, a localized T2-multislice Turbo rapid acquisition with relaxation enhancement (T2 TurboRARE; TE/TR=33ms/2,500ms) was used while a T1-weighted RARE sequence (T1 RARE; TE/TR=6.5ms/1,500ms) was applied for obtaining T1-weighted images. Volumetric analysis of BrMs was performed on MRI DICOM files using a segmentation tool in the ITK-Snap software ^73^.

### Tumor onset and progression

Established BrM was defined as an MRI volume > 0.1 mm^3^ on wildtype, *Entpd1*KO, *Nt5e*KO, *A2ar*KO and *Nt5eA2ar*KO mice. Tumor progression was monitored by weekly MRI measurements. Animals were observed daily and euthanized after showing signs of neurological deficit (lethargy, failure to walk, or loss of > 20% body weight) or reached a maximum brain tumor volume of > 100 mm^3^ based on MRI measurements. BrM-free survival was defined from tumor injection to tumor detection. Symptom-free survival was defined from tumor detection to mouse death.

### Quantitative mass spectrometry

Mouse brains were harvested in an isopentane and dry ice/ethanol bath and stored at -80°C until use. For absolute quantification of endogenous adenosine (ADO), adenosine-D2 calibrants were spotted on control brain sections to build the calibration curve and diaminonaphtalene (DAN, Merck, Cat#56451) matrix mixed to ADO-^13^C_5_ internal standard was sprayed onto 10 µm thick sections and these control brain sections with the automatic TM Sprayer (HTX Technologies, LLC, 4.1) prior analysis. For relative quantification of ATP, DAN-HCl matrix spiked with ATP-^13^C_10_ (Merck, Cat#710695-1MG) internal standard was used and the intensity ratio was calculated between similar regions of interest. Data acquisition at 80 µm spatial resolution was performed using 7T MALDI-FTICR (SolariX XR, Bruker Daltonics, 2.0.78 and 4.1) with Continuous Accumulation of Selected Ions negative mode for adenosine distribution and with full scan negative mode for ATP distribution and subsequently analyzed with Multimaging software (Aliri®, 1.1).

### Liquid-chromatography/mass spectrometry

Mouse BrMs were macrodissected based on MRI images. For healthy control brains, a similar area of the brain was harvested. Human brain metastasis tissue was obtained directly after surgery. Information on the tumor location and tumor type of human brain metastasis is in Supplementary Table 1. Immediately after harvesting, brain tissue was weighted, harvested in a liquid nitrogen bath and stored at -70°C until use. Samples were extracted with 90% ACN solution containing the corresponding isotopically labelled Internal Standards at 5nM concentration (adenosine, ^13^C_10_H_13_^15^N_5_O_4_, AMP ^13^C_10_H_14_^15^N_5_O_7_P, ADP C_10_H_15_^15^N_5_O_10_P_2_, and ATP ^13^C_10_H_16_N_5_O_13_P_3_). Samples were homogenized using a bead beater MP Fast Prep 24 and centrifuged for 15 minutes at 21000 rcf/min. Supernatants were transferred to glass vials for LC-MS analysis. All experimental samples were measured in a randomized manner. Pooled quality control (QC) samples were prepared by mixing equal aliquots from each processed sample. Multiple QCs were injected at the beginning of the analysis in order to equilibrate the LC-MS system. A QC sample was analyzed after every 5^th^ experimental sample to monitor instrument performance throughout the analytical sequence.

LC-MS/MS high-resolution untargeted analysis was carried out in a Vanquish Horizon UHPLC system coupled to an Orbitrap Exploris 240 mass spectrometer (Thermo Scientific, MA, USA) in positive & negative H-ESI (heated-electrospray ionization) modes. Chromatographic separation was carried out on an Atlantis Premier BEH Z-HILIC column at a flow rate of 0.25 mL/min. The mobile phases consisted of water: acetonitrile (9:1, v/v; mobile phase A) and acetonitrile: water (9:1, v/v; mobile phase B), which were modified with a total buffer concentration of 10 mM ammonium acetate. The aqueous portion of each mobile phase was pH-adjusted (pH 9.0 via addition of ammonium hydroxide). The gradient (15 min total run time including re-equilibration) was started at 95% B and held for 2 minutes, ramped to 50% B in 6 minutes, held at 50% B for 2 minutes, and back to initial conditions in 1 minutes. Column temperature was maintained at 40°C, the autosampler was set to 4°C and sample injection volume was 6 µL. Analytes were recorded via a full scan with a mass resolving power of 120,000 over a mass range from 60 – 900 *m/z* (RF lens: 70%). To obtain MS/MS fragment spectra, data-dependent acquisition was carried out (resolving power: 15,000; scan time: 22 ms; stepped collision energies [%]: 30/50/70; cycle time: 900 ms). Ion source parameters were set to the following values: spray voltage: 3500 V / -3000 V, Sheath gas: 30, Auxiliary gas: 5, Sweep gas: 0, ion transfer tube temperature: 350°C, vaporizer temperature: 300°C. Data was processed using Trace Finder 5.1. Sample concentration was estimated using the ratio to the isotopically labeled internal standards. Inosine levels were estimated using Adenosine Internal standard and Inosine Monophosphate using Adenosine Monophosphate IS. Metabolite amounts were normalized to the weight of harvested brain tissue of each sample.

### Whole brain radiotherapy

Whole brain radiotherapy (WBRT) was applied using the Small Animal Radiation Research Platform (SARRP, X-Strahl Ltd) ^26, 27^. Mice were anesthetized with isoflurane (2.5%) and imaged with an on-board Cone Beam computed tomography (CT) operating at 60 kV and 0.8 mA. CT images were loaded in the Muriplan software (X-Strahl, 2.2.1), and individual isocenters were selected for radiotherapy. WBRT was applied as 2 Gy on five consecutive days and a 10 x 10 mm collimator as 1 arc operating at 220 kV and 13 mA with 5.2 cGy/s. WBRT treatment started at d0.

### Pharmacological treatment

The CD39 inhibitor POM-1 (Tocris, Cat#2689) and the ADORA2A inhibitor SCH58261 (Selleckchem, Cat#S8104) were injected daily intraperitoneally at 5 mg/kg and 10 mg/kg, respectively, starting on day 0 after treatment allocation of tumor-bearing mice. *In vivo*Mab anti-mouse CD73 (TY/23, Bio X Cell, Cat#BE0209, RRID: AB_10950310) or *Invivo* Mab rat IgG2 isotype control (clone 2A3, Bio X Cell, Cat#BE0089, RRID:AB 1107769) was dosed at 200 µg/mouse every three days, starting on day 1 after treatment allocation of tumor-bearing mice. *InVivo*Plus anti-mouse PD1 (clone RMP1-14, Bio X Cell, Cat#BP0146, RRID: AB_10949053) or *InVivo*Plus rat IgG2 isotype control (clone 2A3, Bio X Cell, Cat#BP0089, RRID: AB_11077692) was administered intraperitoneally at a dose of 250 µg/mouse every three days, starting on day 0 after treatment allocation of tumor-bearing mice.

### Combination trial

Mice were stratified into different treatment groups based on volumetric analysis of MRI data to achieve group allocations with similar metastatic burden (tumor size, amount of metastatic lesions) at trial initiation. Pharmacological treatments started with the first radiation dose on day 0. For the short-term trials, brains were harvested on day 14 after treatment initiation. BrM burden was evaluated by weekly MRI measurements. Animals were observed daily and euthanized when showed signs of neurological deficit (lethargy, failure to walk, or loss of > 20% body weight) or reached a maximum brain tumor volume of > 100 mm^3^ based on MRI measurements. Area under the curve (AUC) was determined from absolute tumor growth of the individual mice. AUC was normalized to the survival time in days (AUC/days). Mice were pharmacologically treated until the predetermined trial end point or until they developed symptoms from BrM or reached the maximum brain tumor volume (100 mm^3^). For *in vivo* experiments, blinding was not performed as knowledge of the experimental conditions was required during treatment as well as for veterinary and staff monitoring of *in vivo* experiments. For subsequent analysis, automated analysis tools were used and/or the experimenter was blinded to group allocation.

### Multiplexed cytokine assay

BrMs were macrodissected based on MRI images 14 days after treatment start. Protein lysates were isolated using the Bio-Plex Cell Lysis Kit (Bio-Rad, Cat#171304011) and quantified with the Invitrogen Qubit Protein Assay kit (Thermo Fisher Scientific, Cat#Q33211). Multiplexed cytokine analyses were performed with the Bio-Plex Pro Mouse Cytokine 31-plex assay (Bio-Rad, Cat#M60009RDPD) according to manufacturer’s instructions. Raw values were obtained using the Bio-Plex 200 system version 6.1. Cytokine levels were normalized to total protein content of each sample.

### Flow cytometry analysis and cell sorting

BrMs were macrodissected based on MRI images and dissociated using the Brain Tumor Dissociation Kit (Miltenyi Biotec, Cat#130-095-942) and a single cell-suspension was generated using the OctoMACS dissociator (Miltenyi Biotec). Cell suspensions were filtered through a 70µm mesh filter followed by incubation with Myelin Removal Beads II (Miltenyi Biotec, Cat#130-096-433). Cell suspensions were incubated for 15 minutes at 4°C with Mouse BD Fc block (BD Biosciences, Cat#553142, 2.4G2, RRID:AB_394656) followed by incubation with directly conjugated antibody panels for 15 minutes at 4°C. When required, the brilliant stain buffer (BD Biosciences, Cat#566349) was used. Cell suspensions were resuspended in a live-dead staining (Thermo Fisher Scientific, Cat#L34962) or stained with eFluor (Thermo Fischer Scientific, Cat#65-0865-18). Flow cytometry analyses were performed on a BD Fortessa. Flow cytometry analysis sorting was performed on a BD FACS Aria Fusion and cells were sorted directly into TRIzol LS and snap frozen on dry ice. Antibody information is in Supplementary Table 2. Analyses were performed in FlowJo (BD Biosciences, 10.8).

### RNA sequencing and gene expression analysis

Sequencing was performed at Single Cell Discoveries (Utrecht, Netherlands) using an adapted version of the CEL-seq protocol. In brief, total RNA was extracted using a standard TRIzol (Invitrogen) protocol and used for library preparation and sequencing. mRNA was processed as described previously^74, 75^, following an adapted version of the single-cell mRNA seq protocol of CEL-Seq. Samples were barcoded with CEL-seq primers during a reverse transcription and pooled after second strand synthesis. The resulting cDNA was amplified with an overnight *in vitro* transcription reaction. From this amplified RNA, sequencing libraries were prepared with Illumina Truseq small RNA primers. Paired-end sequencing (2x150 bp) was performed on the Illumina HiseqX platform. Read 1 was used to identify the Illumina library index and CEL-Seq sample barcode. Read 2 was aligned to the mm10 RefSeq transcriptome using BWA^76^. Reads that mapped equally well to multiple locations were discarded.

FASTQ files were generated as part of the MapAndGo pipline (https://github.com/anna-alemany/transcriptomics/tree/master/mapandgo), in which sample barcodes whitelist are read, and FASTQ files parsed wherein reads were divided according to their sample barcode. A barcode correction of up to one Hamming distance was applied if necessary. Raw fastq files were quality checked by using the fastqc function of FASTQC package (v. 0.11.9), and multiple fastq files per sample were concatenated by “cat” function. FASTP package (v. 0.23.1) was used for pre-processing of fastq files (adaptor trimming and cutting of low-quality reads). Pre-processed files were then passed to STAR aligner (v. 2.7.3a) for read alignment against mouse genome (GRCm38.p6) using default parameters in addition to quantMode = “GeneCounts” to produce raw read counts.

For further downstream analysis in R (v. 4.2.1) operated in RStudio (v. 1.1.453), HTSeq count files (one per sample) were generated based on HTSeq (Anders et al., 2015)^77^.

Groupwise comparison, differential gene expression and further analysis was performed with BioMart package (v. 2.34.1) and DESeq2 (v. 1.18.1) (Love et al., 2014)^78^. Data for heatmaps and PCA clustering were generated of variance-stabilized transformed data (equals log2 transformation).

For pathway analyses and gene annotation, data were filtered for significantly differentially expressed genes, respecting a basemean greater than 20 and an adjusted p-value (padj) less than 0.05 (= FDR 5%). Pathway analysis and gene annotation were performed with Metascape ^79^ or clusterprofiler ^80^ and enrichplot^81^.

### GeoMx spatial transcriptomics

Paraffin-embedded brain slides were processed according to the GeoMx® DSP slide preparation manual. After being deparaffinized and hydrated by Leica Biosystems BOND RX, the slides were incubated with Whole Transcriptome Atlas probe mix overnight (NanoString, Cat#GMX-RNA-NGS-MsWTA-4). The slides were stained with PANCK (1:50, Novus Biologicals, Cat#NBP2-33200AF594, AE1/AE3, RRID: AB_2924199), IBA1 (1:100, Abcam, Cat#ab221790, EPR16589, RRID: AB_2924254), CD3 (1:100, Bio-Rad, Cat#MCA1477A647, CD3-12, RRID: AB_10845948) and SYTO83 (1:25, ThermoFisher Scientific, Cat#S11364). Regions of interest were demarcated in GeoMx® DSP, and oligos from PANCK^+^ and PANCK^-^ regions were collected upon UV cleavage via a digital micromirror device as different areas of illuminations. The oligos were uniquely indexed using Illumina’s i5xi7 dual-indexing system. PCR reactions were purified and libraries were paired-end sequenced (2x75) on a NovaSeq system (Illumina). 48 areas of illumination were profiled and 19,962 genes passed quality filters. Data was normalized to the third quartile. Principal component analysis (PCA) was used to visualize the dataset in a three-dimensional space. For deconvolution of cell composition by SpatialDecon, cell mixing proportions were estimated using the R code published in NanoString’s Github site^30,31^.

### Visium spatial transcriptomic profiling

Fresh frozen brain tissue was embedded in Tissue-Tek O.C.T. Compound (12351753, Fisher Scientific). RNA was isolated from serial sectioned tissues totalling 80 µm thickness and RNA integrity number was calculated using the Agilent 4200 TapeStation system. Frozen tissue sections at 10 µm were cut in the cryostat for spatial transcriptomic profiling. Spatial transcriptomics sequencing libraries were prepared with 10x Genomics Visium Spatial Gene Expression Slides and Reagent Kit according to manufacturer’s instructions. The samples were loaded on a NovaSeq 6000 System (Illumina) using NovaSeq 6000 SP Reagent Kit v1.5 (500 cycles, 20028402, Illumina). Raw sequencing data were processed using the 10x Genomics Space Ranger pipeline and mapped to GRCm38 mouse reference genome (mouse reference mm10-2020-A). Seurat was used to perform pre-processing and spatial clustering of spots based on transcriptome profiles. CellChat functions for spatial transcriptomics data were used to infer spatial cell-cell interactions^82, 83^.

### Single cell RNA sequencing

Brain was harvested from tumor-bearing mice of n=3/group for FACS sorting. Sorted cells were positive for CD45 and negative for EPCAM, cleaved caspase 3 and propidium iodide. Cells were further processed according to the standard 10 x Genomics workflow. Libraries were generated using the Chromium Next GEM Single Cell 3’ v3.1 kit (10x genomics, Cat#PN-1000147). The Bioanalyzer High Sensitivity DNA chip (Agilent) and Qubit dsDNA High Sensitivity Kit (Thermo Fisher Scientific, Cat#Q32851) were used for cDNA QC and DNA quantification. The sequencing was performed on NovaSeq6000 platform (Illumina) with a sequencing depth of at least 20,000 reads per cell using the NextSeq500/550 high output kit v2.5 (75 cycles; Illumina; Cat#20024906). Fastq files were processed using cellRanger v7.0 using cellranger count pipeline with default arguments. Cellranger output was then used for downstream analysis. The transcriptomes were mapped to GRCm38 mouse reference genome (mouse reference mm10-2020-A). Cells with at least 1,000 UMIs per cell and less than 20% mitochondrial gene content were retained for analysis. The downstream analysis was performed using a combination of different mathematical methods: principal component analysis^84^, Louvain clustering^85^, UMAP^86^. The most variable genes in the dataset were identified and the top 10% were used for dimensionality reduction and clustering. The top differentially expressed genes per cluster were used to identify cell types. In this workflow we used a combination of R (version 4.3.1) and packages scater (version 1.28.0)^87^, scran (version 1.28.2)^88^, ggplot2 (version 3.4.3)^89^. CellChat was used to infer cell-to-cell communication analysis ^37^.

### Tissue preparation and immunostaining

Paraffin-embedded patient sections were processed using a Leica Bond Max automated staining device followed by incubation with primary antibodies: ENTPD1 (1:1000, Atlas Antibodies, Cat#HPA014067, RRID: AB_1848178), NT5E (1:200, Cell Signaling Technologies, Cat#13160, D7F9A, RRID: AB_2716625), ADORA2A (1:200, Atlas Antibodies, Cat#HPA075997, RRID: AB_2732298), ADORA2B (1:15000, Atlas Antibodies, Cat#NBP2-41312, RRID: AB_3073937), ADA (1:10000, Atlas Antibodies, Cat#HPA023884, RRID: AB_1844573) and ADK (1:2000, Atlas Antibodies, Cat#HPA038409, RRID: AB_10675798), followed by incubation with HRP-conjugated secondary antibodies and diaminobenzidine conversion. Haematoxylin and eosin staining were performed on an automated staining device (Leica Autostainer XL). Histopathological scoring of CD39, CD73, ADORA2A, ADORA2B, ADA and ADK expression was obtained from n=94 of non-small cell lung cancer, n=16 of small cell lung cancer, n=48 of breast cancer, n=19 of melanoma, n=20 of colon cancer, n=13 of kidney cancer, n=6 of carcinoma metastasis, n=28 of not specified samples/other tumors and followed the histological score (H score). The H score was calculated as: (0 x % non-stained cells) + (1 x % weak stained cells) + (2 x % moderate stained cells) + (3 x % strong stained cells) using the immunohistochemistry image and the values ranged from 0 to 300. Histological scoring of the samples was blinded to patient’s information.

Similarly, formalin fixed paraffin-embedded (FFPE)-mouse tissue sections were stained for CD3 (1:2800, Abcam, Cat#ab215212, EPR20752, RRID: AB_215212), CD4 (1:200, Cell Signaling Technologies, Cat#252229, D7D2Z, RRID: AB_3073941), CD8a (1:500, Cell Signaling Technologies, Cat#98941, D4W2Z, RRID: AB_2756376) and IBA1 (1:1000, Novus Biologicals, Cat#NBP2-19019, RRID: AB_3073939). Quantification of cell populations was performed with Aperio ImageScope (v12.4.0.50.43) using either a nuclear counting algorithm or a pixel counting algorithm depending on the cell shape.

FFPE-mouse tissue sections were blocked in 3% BSA + 0.1% Triton X-100 in PBS for 1 h at room temperature (RT), followed by incubation with primary antibodies against EPCAM (1:1000, Abcam, Cat#ab71916, RRID: AB_1603782), IBA1 (1:500, Wako Chemicals, Cat#019-19741, RRID: AB_839504), and TMEM119 (1:500, Synaptic Systems, Cat#400004 RRID: AB_2744645) in 1.5% BSA overnight at 4°C. Fluorophore-conjugated secondary antibodies were used at a dilution of 1:500 in 1.5% BSA in PBS for 1 h at RT. Images were taken using the Yokogawa CQ1 software (SM 80J01A01-01E, 1.04.04.02).

For multiplexed histology, mouse FFPE sections were stained using the Opal Polaris 7 Color Kit (Akoya Biosciences Inc., Cat#NEL861001KT) based on tyramide signal amplification immunostaining technique. Nuclei detection was performed using Spectral DAPI (Akoya Biosciences, Inc., Cat#FP1490). 7-plex stainings were performed on a LabSat^TM^ Research Automated staining instrument (Lunaphore Technologies SA) targeting CD4 (1:450, Abcam, Cat#ab183685, EPR19514, RRID: AB_2686917), CD8a (1:450, Cell Signaling, Cat#98941S, D4W2Z, RRID: AB_2756376), EPCAM (1:180. Abcam, Cat#ab71916, RRID: AB_1603782) and IBA1 (1:250, Wako Chemicals, Cat#019-19741, RRID: AB_839504). Multiplex stainings were acquired on Vectra Polaris (Akoya Biosciences, Inc.) using MOTiF^TM^ technology which provided unmixed whole slide scans of complete tumors at 0,5 µm/pixel. Multispectral image analysis was performed with Phenochart^®^ version 1.0.12, InForm^®^ Image Analysis Software (Akoya Biosciences, Inc.) and HALO^TM^ software (Indica Labs).

### Quantitative PCR with reverse transcription analysis

RNA was isolated using TRIzol, DNase treated. RNA was transcribed to cDNA using the High-Capacity cDNA Reverse Transcription Kit (Thermo Fisher Scientific, Cat#4368814). Assays were run in triplicate following the instructions of the TaqMan Gene Expression Master Mix (Thermo Fisher Scientific, Cat#4369016), and expression of *Entpd1* (Mm00515457_m1), *Nt5e* (Mm00501912_m1), *A2ar* (Mm00802075_m1), *A2br* (Mm00839292_m1), *Ada*

(Mm_00545720_m1) and *Adk* (Mm00612772_m1) was normalized to ubiquitin C (Thermo Fisher Scientific, Mm99999915_g1) and glyceraldehyde 3-phospahte dehydrogenase (Thermo Fisher Scientific, Mm02525934_g1) for each sample.

### Measurement of AMPase enzymatic activity

AMPase activity was measured using the Malachite Green Phosphate Detection Kit (R&D Systems, Cat#DY996). 10x10^3^ 99LN, scrambled-99LN, *Nt5e*-deficient cells were seeded in 96-well-plates. 48h later cells were washed at RT with orthophosphate-free reaction buffer (20 mM HEPES, 5,6 mM D-glucose, 5 mM KCl, 1 mM MgCl_2_, 100 mM NaCl, 1,8 mM CaCl_2_, pH 7-8) followed by incubation with 1 mM AMP (Thermo Fisher Scientific, Cat#11986851) for 40 minutes at 37°C. Cells without nucleotides and wells without cells were incubated as well. The reaction was stopped by adding pre-chilled 10% Trichloroacetic acid (Sigma, Cat#T9159). The supernatants were then transferred to a new plate and diluted 1:5 with orthophosphate-free reaction buffer. Per 50µl sample, 30µl Malachite Green Reagent A were added, followed by addition of 100µl distilled water and 30µl Malachite Green Reagent B. After 40 minutes incubation, the mixture was measured by monitoring the absorbance at 630 nm with a microplate reader. µmol phosphate concentrations were estimated from a phosphate standard curve.

### Cell viability assay

Cell growth rate was determined by MTS cell viability assay (Promega, Cat#G3582). Cells were plated in triplicate at 1 x 10^3^ cells in 96-well-plates, with media changes every 48 hours.

### RNAseq expression data from clinical BrM samples

RNAseq raw count data of sorted cell populations from healthy donors and human brain metastases (published in^20, 24^) were downloaded from the Brain TIME website (https://joycelab.shinyapps.io/braintime/).

### Visualization of human spatial transcriptomics

Spatial transcriptomics data from a Sudmeier et al., 2022 ^23^ was downloaded from GEO. After quality control, the per-spot expression values were library-size normalized and the gene expression was visualized using the R package ggplot2.

### Quantification and statistical analysis

Summary data are presented as mean ± standard error of the mean (sem), individual data points with line at mean, box and whisker plot indicating the median or violin plots with lines indicating min, max and median using GraphPad Prism software v9 or R/ggplot2. Data was checked for normal distribution. Statistical analyses were performed with GraphPad Prism software v9 or R v4.2.1 using the indicated statistical tests (two-tailed), with *P*<0.05 considered statistically significant as indicated in each figure legend. When significant, the exact *P* values are reported in each figure. Further information on the statistical test and number of biological replicates (n) used for every assay is found in the corresponding figure legend.

---

**Extended Data Figure 1.**
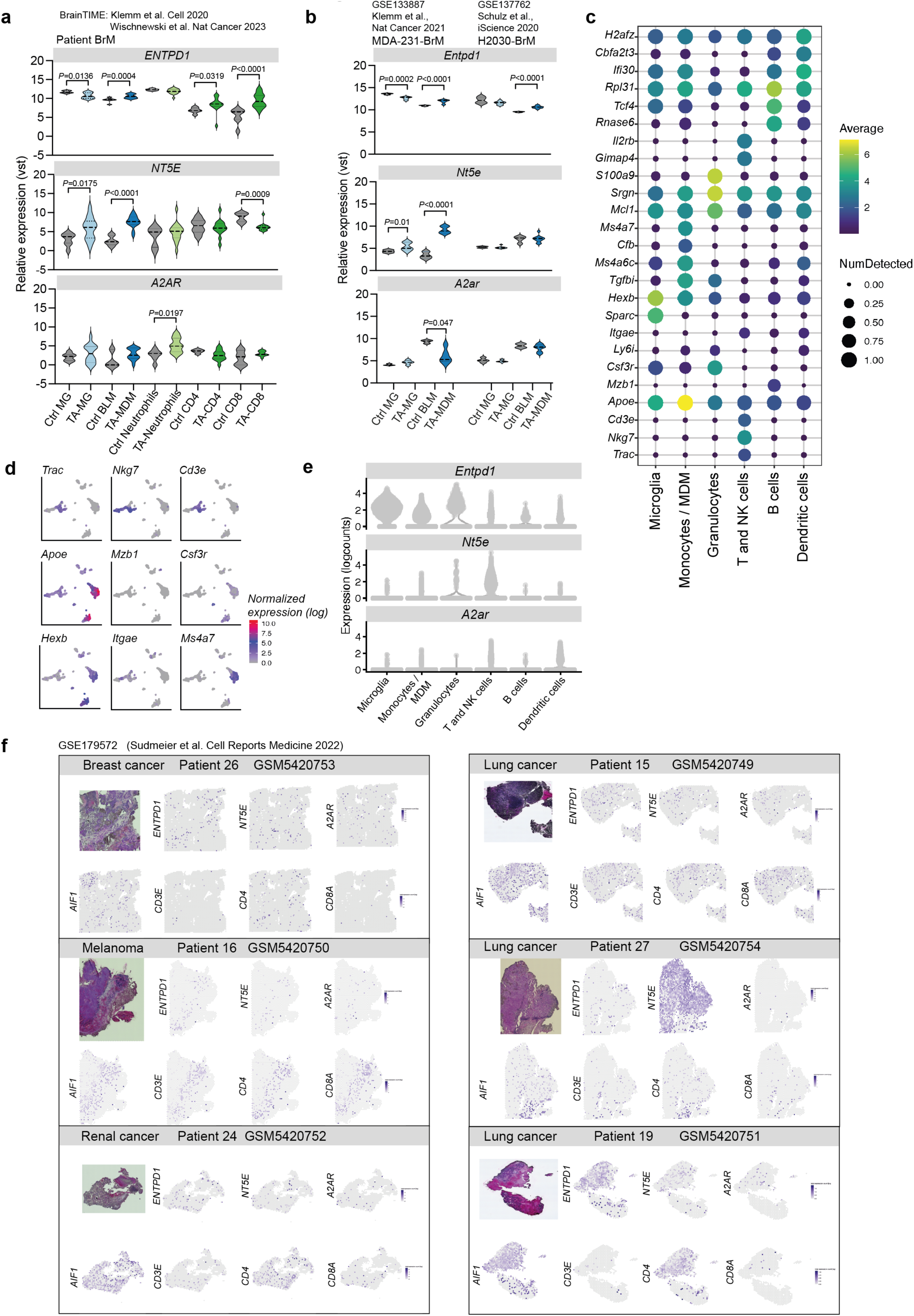
ATP-adenosine signaling components in brain metastasis. **a**, Quantification of gene expression levels of indicated factors based on bulk RNA seq of human FACS purified cell populations. Data were obtained from Klemm et al., 2020 BrainTIME (Ref20) and Wischnewski et al., 2023 (Ref24). Unpaired t-test was used. **b**, Gene expression data of purine and adenosine/inosine signaling components in control and tumor-associated cells in murine breast-to-brain MDA experimental BrM and lung-to-brain H2030 BrM. Data were obtained from Schulz et al. 2020 (Ref22, and Klemm et al., 2021 (Ref21). **c**, Discriminative marker genes for each functionally enriched state detected by scRNAseq of breast-to-brain 99LN-BrM. **d**, UMAP of scRNAseq from sorted CD45+ cells of 99LN-BrM showing the indicated genes. **e**, Expression of the indicated factors in the different cell types in 99LN-BrM model analyzed by scRNAseq. **f**, Spatial gene expression profiles for the indicated genes in human BrM samples. Data were obtained from Sudmeier et al., 2022 (Ref23). Salamero-Boix et al. Extended Data Figure 2

**Extended Data Figure 2.**
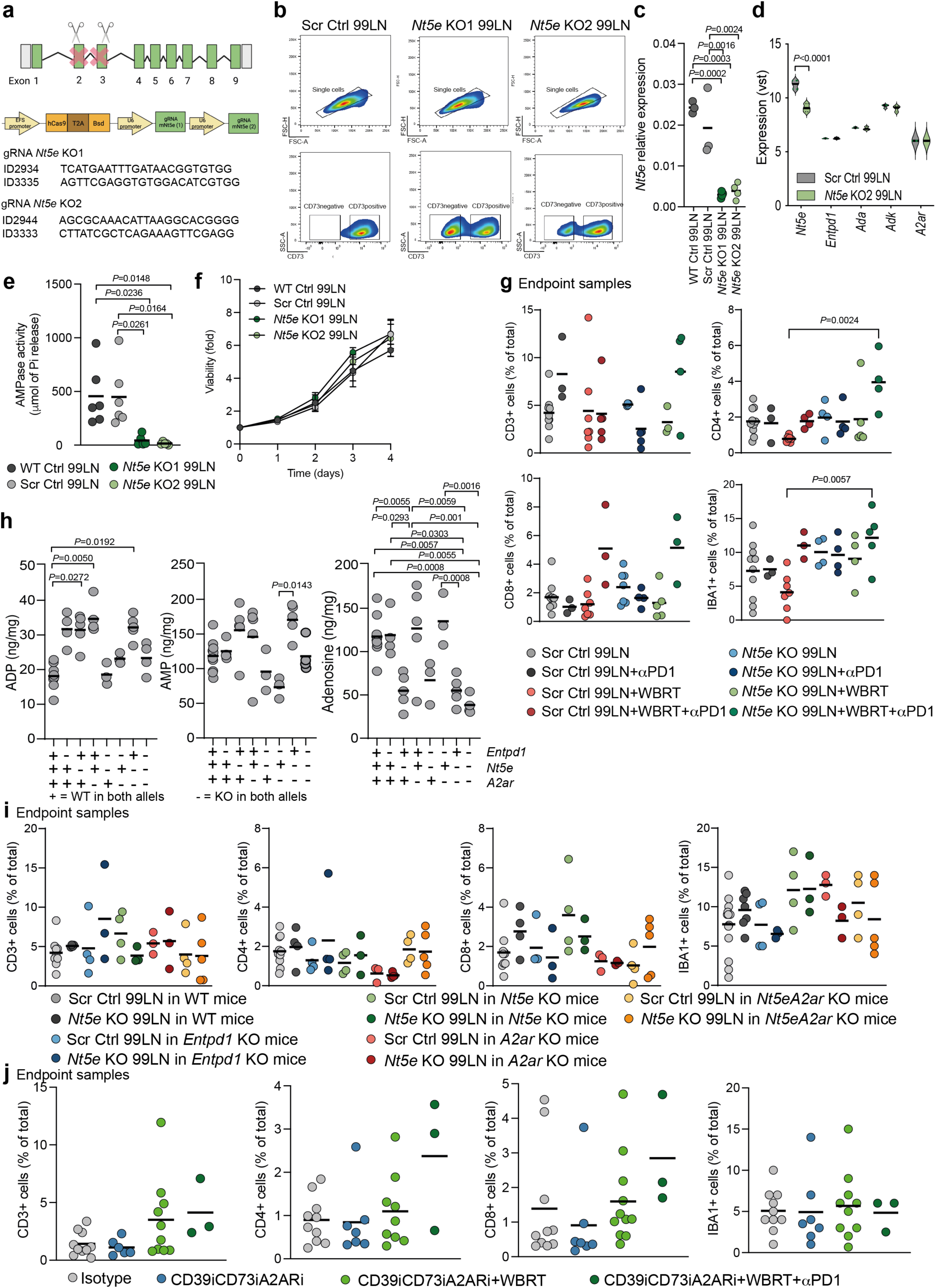
Manipulation of *Entpd1*, *Nt5e* and *A2ar* in murine 99LN-BrM breast-to-brain metastasis. **a**, Scheme of the targeting strategy used to generate the *Nt5e*-deficient tumor cell line. **b**, Flow plots of *Nt5e*-proficient and *Nt5e*-deficient cells upon *Nt5e* genetic targeting in the 99LN tumor cell line. **c**, *Nt5e* relative expression in the indicated tumor cell lines, n=3. **d**, Expression of the indicated factors in the indicated tumor cell lines, n=3. padj was obtained by Wald test and corrected for multiple testing using the Benjamini and Hochberg method. **e**, AMPase activity of the indicated tumor cell lines, n=6 Ordinary one-Way ANOVA with Tukey’s multiple comparisons test was used. **f**, Measurement of cell viability in the indicated cell lines using MTS assay (n=4). Ordinary one-way ANOVA with Tukey’s multiple comparisons test was used. **g**, Quantification of CD3+, CD4+, CD8+ and IBA1+ cells in sections of 99LN-BrM from Scr Ctrl 99LN (n=10), Scr Ctrl 99LN+αPD1 (n=3), Scr Ctrl 99LN+WBRT (n=7), Scr Ctrl 99LN+WBRT+αPD1 (n=5), *Nt5e*KO 99LN (n=4), *Nt5e*KO 99LN+ αPD1 (n=4), *Nt5e*KO 99LN+WBRT (n=5), *Nt5e*KO 99LN+WBRT+αPD1 (n=4). Kruskal-Wallis test with Dunn’s multiple comparisons test was used for CD3, CD4 and CD8. Ordinary one-Way ANOVA test with Tukey’s multiple comparisons test was used for IBA1. **h**, Quantification of ADP, AMP and adenosine in the indicated mouse strains: Wildtype (n=10), *Entpd1*KO (n=4), *Nt5e*KO (n=5), *A2ar*KO (n=5), *Entpd1Nt5e*KO (n=4), *Entpd1A2ar*KO (n=3), *Nt5eA2ar*KO (n=5) and *Entpd1Nt5eA2Ar*KO (n=4). Kruskal-Wallis test with Dunn’s multiple comparisons test was used for ADP and AMP. Ordinary one-Way ANOVA test with Tukey’s multiple comparisons test was used for adenosine. **i**, Quantification of CD3+, CD4+, CD8+ and IBA1+ cells in sections of BrM from Scr Ctrl 99LN in WT mice (n=12), *Nt5e*KO 99LN in WT mice (n=4), Scr Ctrl 99LN in *Entpd1*KO mice (n=4), *Nt5e*KO 99LN in *Entpd1*KO mice (n=4), Scr Ctrl 99LN in *Nt5e*KO mice (n=4), *Nt5e*KO 99LN in *Nt5e*KO mice (n=3), Scr Ctrl 99LN in *A2ar*KO mice (n=3) *Nt5e*KO 99LN in *A2ar*KO mice (n=3), Scr Ctrl 99LN in *Nt5eA2ar*KO mice (n=4), *Nt5e*KO 99LN in *Nt5eA2ar*KO mice (n=5). Ordinary one-way ANOVA with Tukey’s multiple comparisons test was used for CD3. Kruskal-Wallis test with Dunn’s multiple comparisons test was used for CD4, CD8 and IBA1. **j**, Quantification of CD3+, CD4+, CD8+ and IBA1+ cells in sections of BrM from isotype (n=10), CD39iCD73iA2ARi (n=7), CD39iCD73iA2ARi+WBRT (n=8), CD39iCD73iA2ARi+WBRT+αPD1 (n=3). Ordinary one-way ANOVA with Tukey’s multiple comparisons test was used.

**Extended Data Figure 3.**
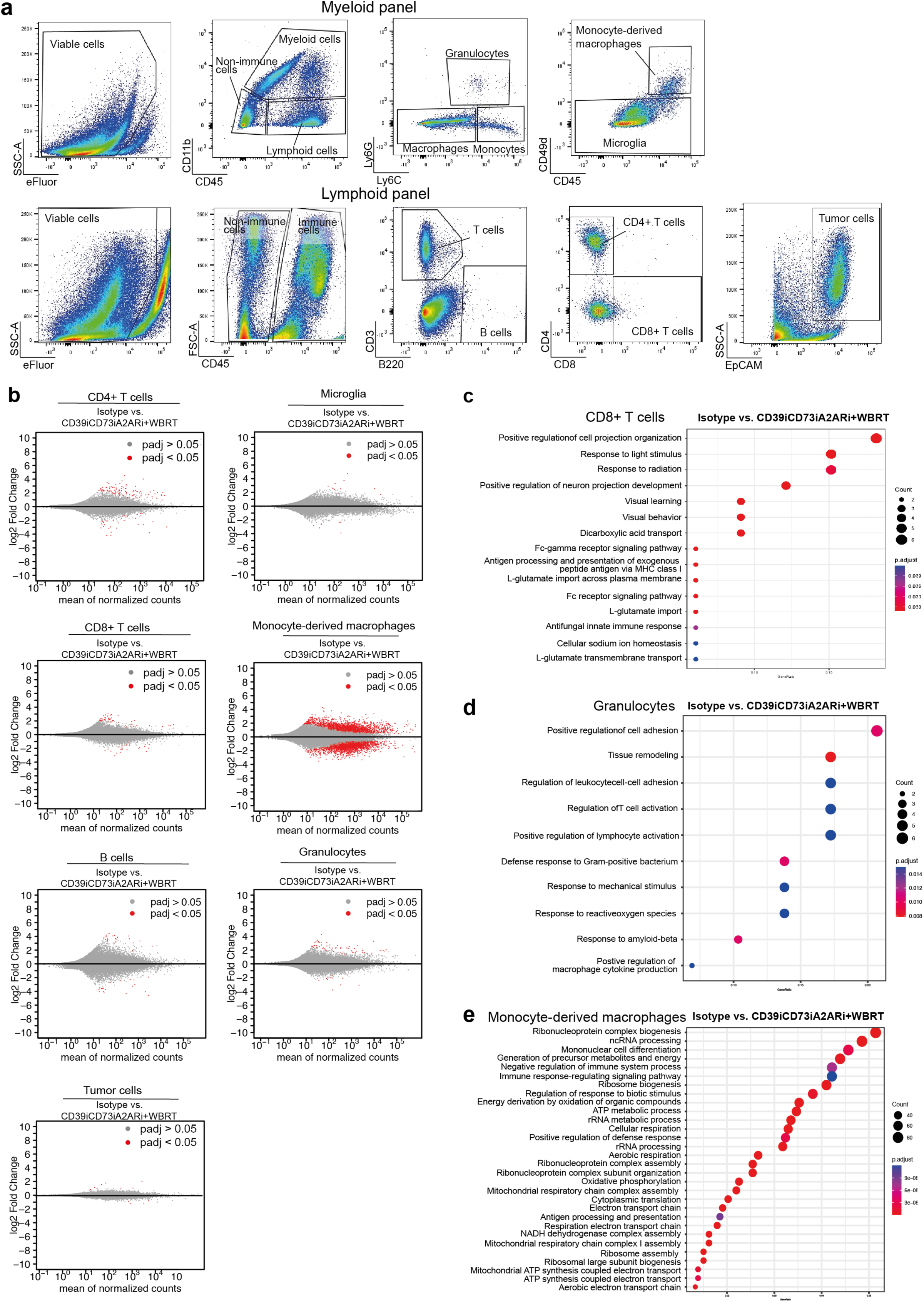
CD39iCD73iA2ARi+WBRT-immune mediated effects in 99LN-BrM. **a**, Gating schemes for flow cytometry analysis of the myeloid and lymphoid compartment of 99LN-BrM. **b**, MA plots for sorted CD4, CD8, B220, microglia, MDM and granulocytes and tumor cells from isotype-treated vs. CD39iCD73iA2ARi+WBRT-treated samples. **c**, Functional annotation of DEG (cut off padj < 0.05) on CD8 sorted T cells from isotype-treated vs. CD39iCD73iA2ARi+WBRT-treated 99LN-BrM samples. Color gradient of red to purple indicates increasing padj; circle size indicates number of genes in pathway. **d**, Functional annotation of DEG (cut off padj < 0.05) on granulocytes sorted cells from isotype-treated vs. CD39iCD73iA2ARi+WBRT-treated 99LN-BrM samples. Color gradient of red to purple indicates increasing padj; circle size indicates number of genes in pathway. **e**, Functional annotation of DEG (cut off padj < 0.05) on monocyte-derived macrophages (MDM) sorted cells from isotype-treated vs. CD39iCD73iA2ARi+WBRT-treated 99LN-BrM samples. Color gradient of red to purple indicates increasing padj; circle size indicates number of genes in pathway.

**Extended Data Figure 4.**
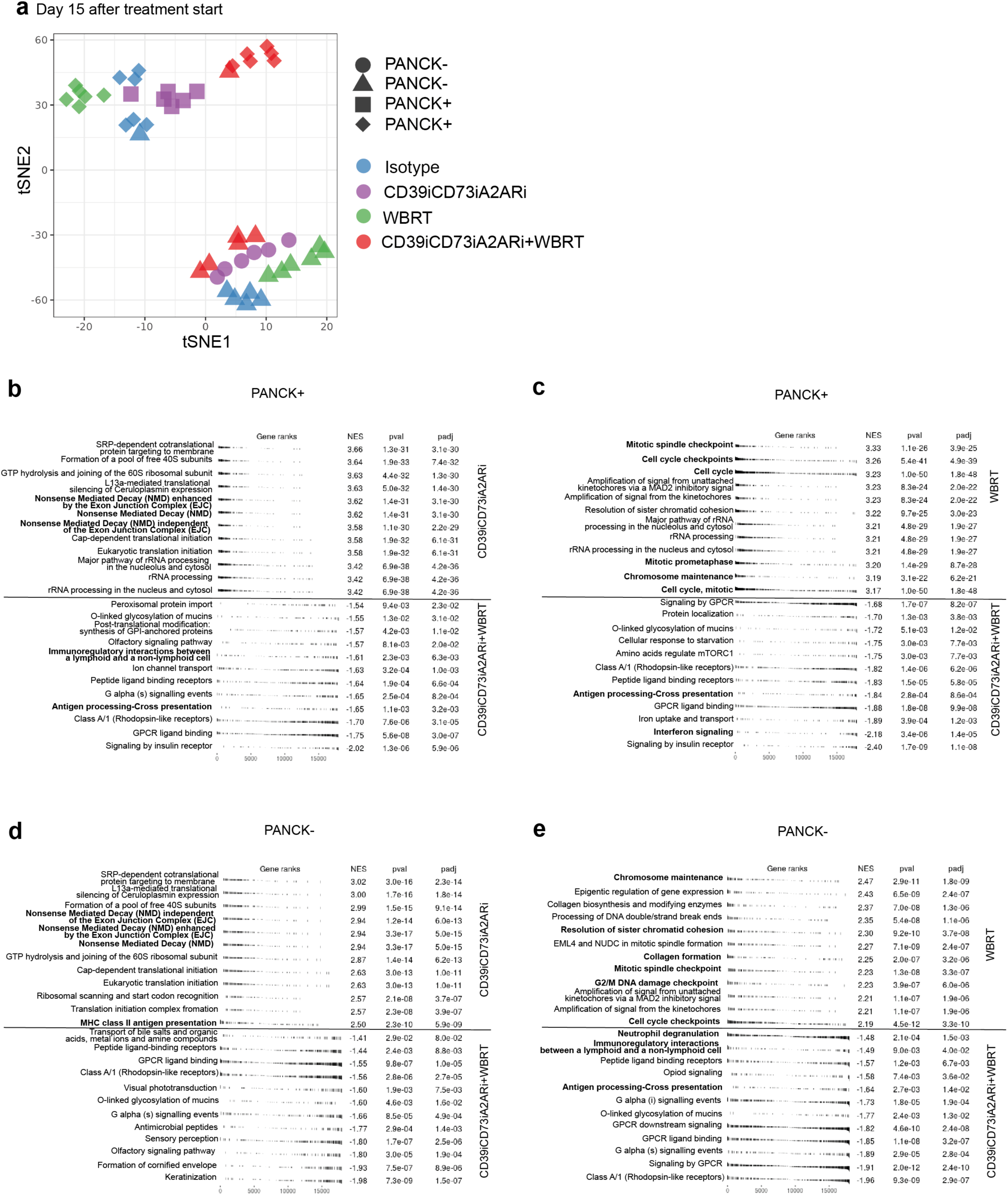
Spatial specificity of gene expression upon CD39/CD73/ADORA2A pharmacological modulation and WBRT in 99LN-BrM. **a**, tSNE representation of PANCK-positive and PANCK-negative areas of the GeoMx analysis in the indicated conditions on 99LN-BrM samples. **b**, Twenty pathways with lowest Padj from the MSigDB GO collection identified by gene set ORA in the DEG set CD39iCD73iA2ARi-treated versus CD39iCD73iA2ARi+WBRT 99LN-BrM for PANCK+ areas. **c**, Twenty pathways with lowest Padj from the MSigDB GO collection identified by gene set ORA in the DEG set WBRT-treated versus CD39iCD73iA2ARi+WBRT 99LN-BrM for PANCK+ areas. **d**, Twenty pathways with lowest Padj from the MSigDB GO collection identified by gene set ORA in the DEG set CD39iCD73iA2ARi-treated versus CD39iCD73iA2ARi+WBRT 99LN-BrM for PANCK-areas. **e**, Twenty pathways with lowest Padj from the MSigDB GO collection identified by gene set ORA in the DEG set WBRT-treated versus CD39iCD73iA2ARi+WBRT 99LN-BrM for PANCK-areas.

**Extended Data Figure 5.**
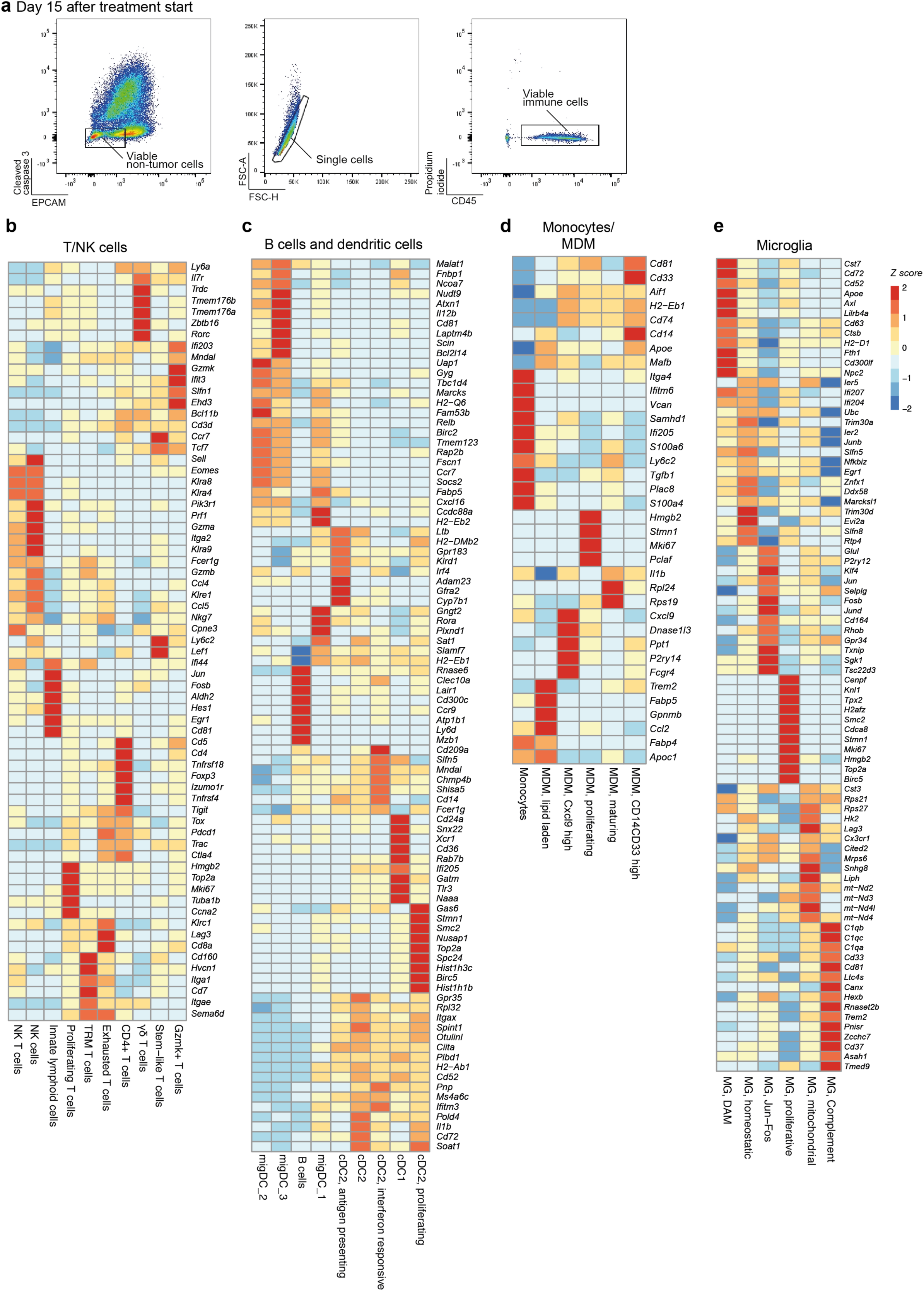
Description of the enriched states detected by scRNAseq of CD45+ cells in 99LN-BrM. **a**, Gating schemes for flow cytometry sorting of CD45+ cells subjected to scRNAseq. n=3/condition pooled for cell sorting. **b,** Heatmap representing the discriminative marker genes for each functionally enriched state detected by scRNAseq for the T/NK cell compartment. **c**, Heatmap representing the discriminative marker genes for each functionally enriched state detected by scRNAseq for the B/dendritic cell compartment. **d**, Heatmap representing the discriminative marker genes for each functionally enriched state detected by scRNAseq for the monocyte/MDM cell compartment. **e**, Heatmap representing the discriminative marker genes for each functionally enriched state detected by scRNAseq for the microglia compartment.

**Extended Data Figure 6.**
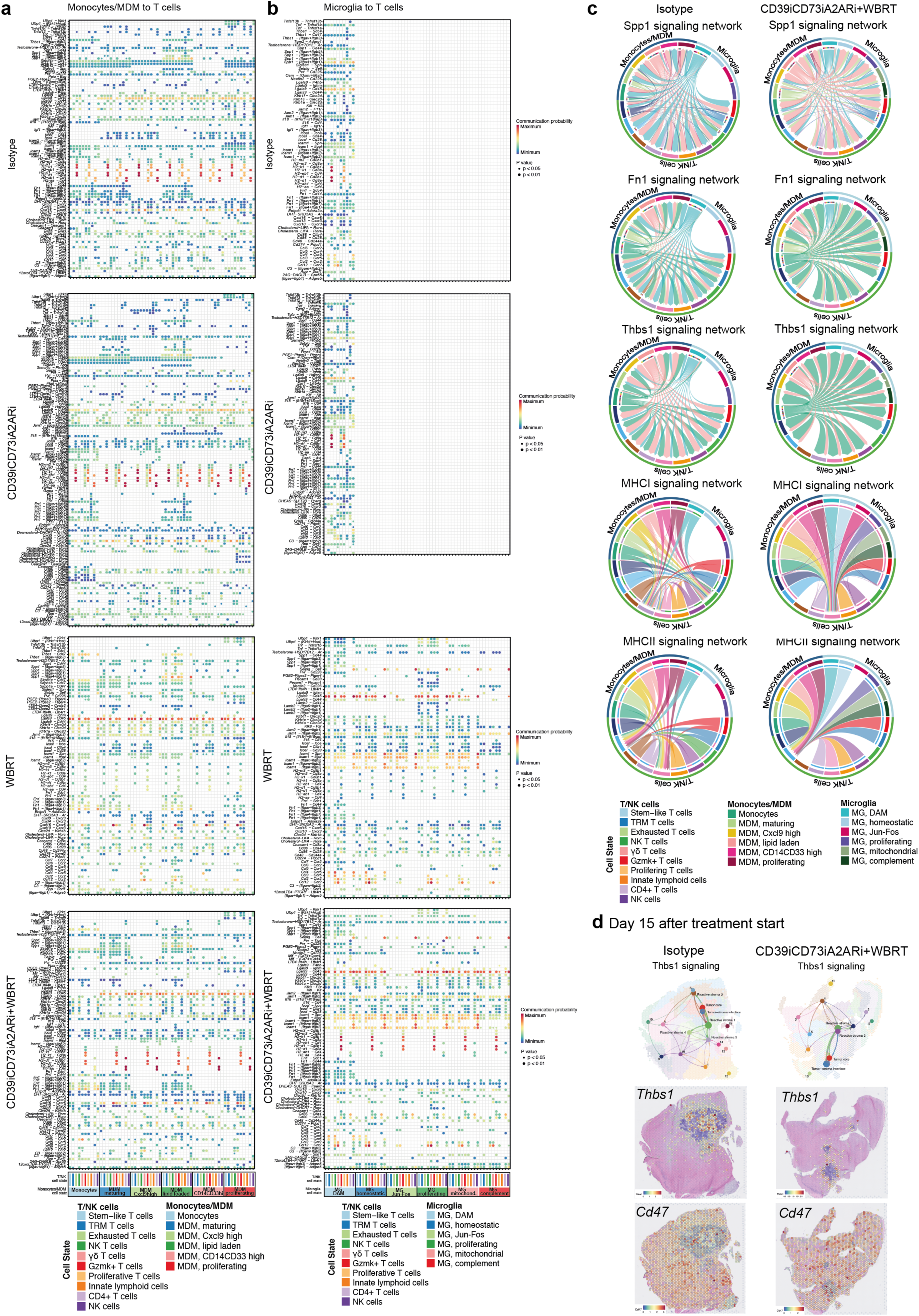
Analysis of ligand-receptor cell interactions between the monocytes/MDM and microglia compartments with the T/NK cell compartment in 99LN-BrM. **a**, Bubble plot depicting in the y axis the most detected ligand-receptor pair interactions and in the x axis the interacting monocytes/MDM and T/NK cell subclusters upon isotype, CD39iCD73iA2ARi, WBRT and CD39iCD73iA2ARi+WBRT treatment. Color gradient of blue to red shows the communication probability. Increased circle size indicates decreased *P* value. **b**, Bubble plot depicting in the y axis the most detected ligand-receptor pair interactions and in the x axis the interacting microglia and T/NK cell subclusters upon isotype, CD39iCD73iA2ARi, WBRT and CD39iCD73iA2ARi+WBRT treatment. Color gradient of blue to red shows the communication probability. Increased circle size indicates decreased *P* value. **c**, Chord diagram showing the cell-cell interactions in isotype and CD39iCD73iA2ARi+WBRT. The perimeter of the chord diagram is color-coded by cellular subtype (colored outer arcs). The inner connecting lines show the interactions between target and receiver cell types. The connecting lines are color coded consistently with cellular subtypes according to the sending cell type. The thickness of each edge (edge weight) is proportional to the interaction strength. **d**, Representation of Thbs1 signaling, Thbs1 and Cd47 expressed as log2 (normalized counts) in isotype and CD39iCD73iA2ARi+WBRT samples.

**Extended Data Figure 7.**
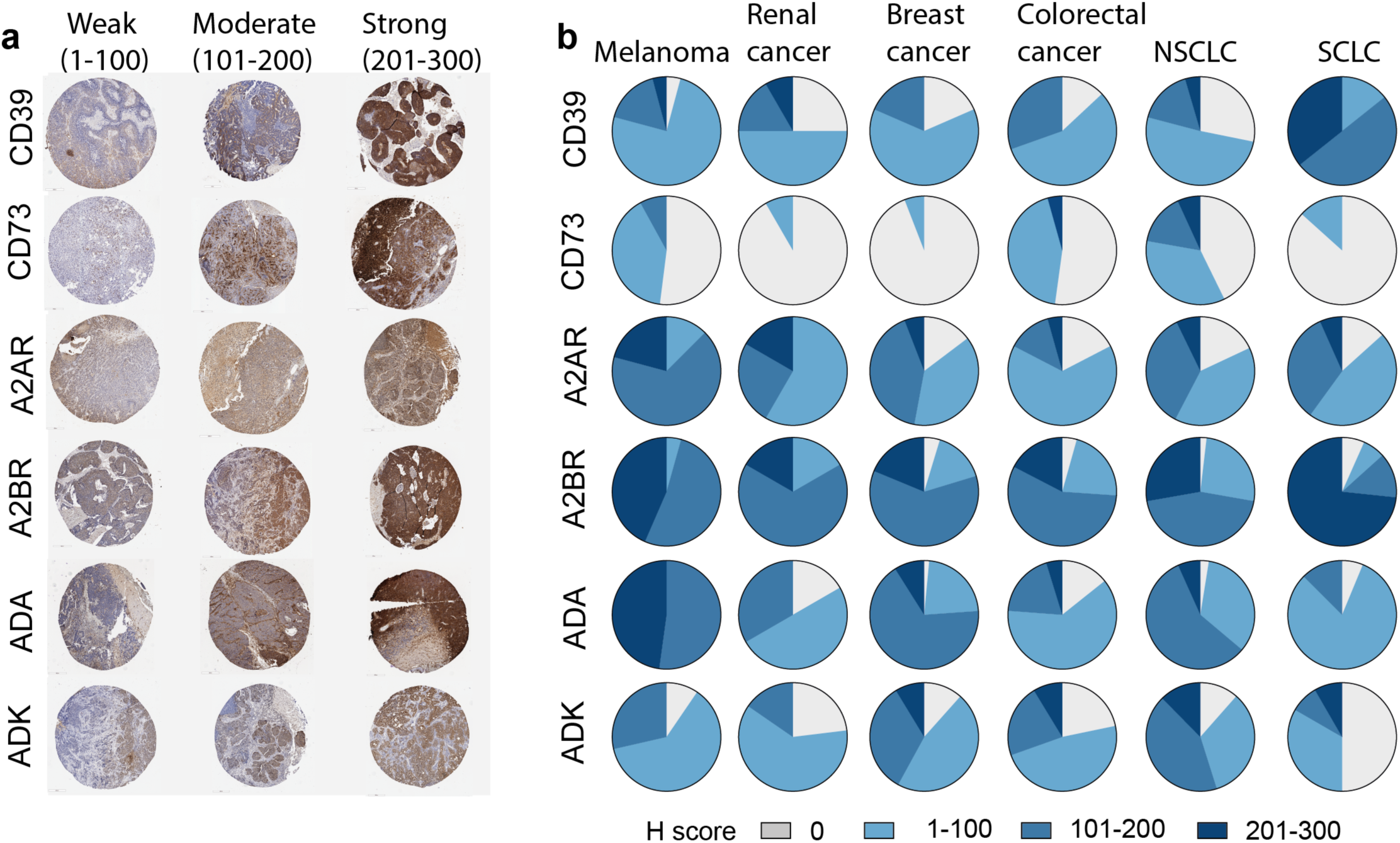
Assessment of ATP-adenosine components in human brain metastasis and adenosine pathway axis targeting in murine lung and melanoma BrM models. **a,** Representative immunohistochemistry images for CD39, CD73, A2AR, A2BR, ADA and ADK belonging to the different staining groups. **b,** Quantification of the tumor histology score (H score) on BrM tissue microarrays according to the indicated tumor types.

## Notes

### Competing Interest Statement

The authors have declared no competing interest.

